# Deep brain stimulation induces white matter remodeling and functional changes to brain-wide networks

**DOI:** 10.1101/2024.06.13.598710

**Authors:** Satoka H. Fujimoto, Atsushi Fujimoto, Catherine Elorette, Adela Seltzer, Emma Andraka, Gaurav Verma, William GM Janssen, Lazar Fleysher, Davide Folloni, Ki Sueng Choi, Brian E. Russ, Helen S. Mayberg, Peter H. Rudebeck

## Abstract

Deep brain stimulation (DBS) is an emerging therapeutic option for treatment resistant neurological and psychiatric disorders, most notably depression. Despite this, little is known about the anatomical and functional mechanisms that underlie this therapy. Here we targeted stimulation to the white matter adjacent to the subcallosal anterior cingulate cortex (SCC-DBS) in macaques, modeling the location in the brain proven effective for depression. We demonstrate that SCC-DBS has a selective effect on white matter macro- and micro-structure in the cingulum bundle distant to where stimulation was delivered. SCC-DBS also decreased functional connectivity between subcallosal and posterior cingulate cortex, two areas linked by the cingulum bundle and implicated in depression. Our data reveal that white matter remodeling as well as functional effects contribute to DBS’s therapeutic efficacy.

## Main text

Despite advances in available treatment options, many people with psychiatric and neurological disorders do not respond to pharmacological, behavioral, or physical therapies (*1*). In these cases, deep brain stimulation (DBS) is increasingly being used to reduce symptoms (*2*). First developed for Parkinson’s disease, this neuromodulation therapy is now being adapted for other neurodegenerative and psychiatric disorders, particularly depression (*3–5*). Importantly, unlike the approach of targeting the gray matter in the subthalamic nucleus, or parts of the motor thalamus, to alleviate movement control deficits (*6–10*), deep brain stimulation for depression instead targets white matter pathways in different parts of the brain (*11–14*). In particular, DBS delivered to the white matter adjacent to subcallosal anterior cingulate cortex (SCC-DBS) (*15–19*) targets the confluence of three white matter tracts: the cingulum bundle, the forceps minor, and the uncinate fasciculus (*20–23*). Delivering DBS to this target has led to sustained long-term symptom reduction in approximately 60-75% of patients that were previously treatment resistant (*24*, *25*). Despite the clinical efficacy of SCC-DBS for depression (*26*), the neurobiological mechanisms engaged by this treatment are still not fully established (*14*). Revealing the therapeutic mechanisms of DBS has the potential to provide improved and new therapeutic targets as well as insight into the underlying pathophysiology of depression.

It has been theorized that SCC-DBS directed to white matter has its beneficial effects by rapidly normalizing activity and functional communication across distributed brain circuits (*3*, *27–32*). While there is evidence to support this functional account of the therapeutic effects (*29–31*, *33*, *34*), the available data are still limited due to the challenges with imaging patients with implanted DBS devices. Often overlooked is the possibility that in addition to its rapid functional effects on neural activity, DBS may promote changes to white matter microstructure concurrently. This idea is supported by the findings that DBS-mediated improvement in mood often takes weeks to fully appear (*3*, *28*); discontinuation of DBS after stable clinical response is generally associated with a slow relapse of symptoms over days to weeks rather than hours (*17*, *22*) and there is a relationship between the degree of white matter integrity prior to starting treatment and the extent of improvement in depression ratings following SCC-DBS (*32*). If DBS causes anatomical remodeling of white matter tracts, it would reveal a previously underappreciated mechanism of therapeutic action and a role for white matter dysfunction in psychiatric disease in general and particularly in depression.

## Results

### SCC-DBS induces selective remodeling of white matter

Matching the location in the brain that has been used for treatment resistant depression, we targeted mini-DBS electrodes to the confluence of the cingulum bundle (CB), uncinate fasciculus and forceps minor in two macaque monkeys (**Figs. 1A** and **B**). DBS leads were implanted unilaterally, allowing for the other hemisphere to act as an internal control to investigate the direct effects of white matter stimulation compared to indirect cross-hemisphere effects. Post-surgery computed tomography (CT) confirmed the DBS lead locations relative to the confluence of the three white matter tracts and this was used to determine the active contact for stimulation (**Fig. 1B**, bottom left). Following a four week period after implantation during which no stimulation was delivered, chronic stimulation (130Hz, 5mA, 90μsec) was delivered for six weeks (**Fig. 1A**). This stimulation regime was used as it mirrors the approach taken in human patients (*3*, *22*, *28*, *30*). Further, it was used because on average the rate of change in depression rating symptoms asymptotes after six weeks stimulation (*3*, *16*, *22*). After lead implantation, including time points both before and after stimulation was started, we assessed the animals’ homecage behavior to see if there were any changes in movement or foraging behaviors. Both the time spent moving and foraging increased after SCC-DBS (one-way repeated measures ANOVA, movement: F_(1,240)_ = 33.85, p < 0.001, foraging: F_(1,240)_ = 17.26, p < 0.001, **Fig. S1**). No neurological deficits were observed after SCC-DBS. Thus, despite both animals being healthy and stimulation being unilateral, SCC-DBS induced a slight change in behavior confirming that it was having an effect on the brain.

**Figure 1:**
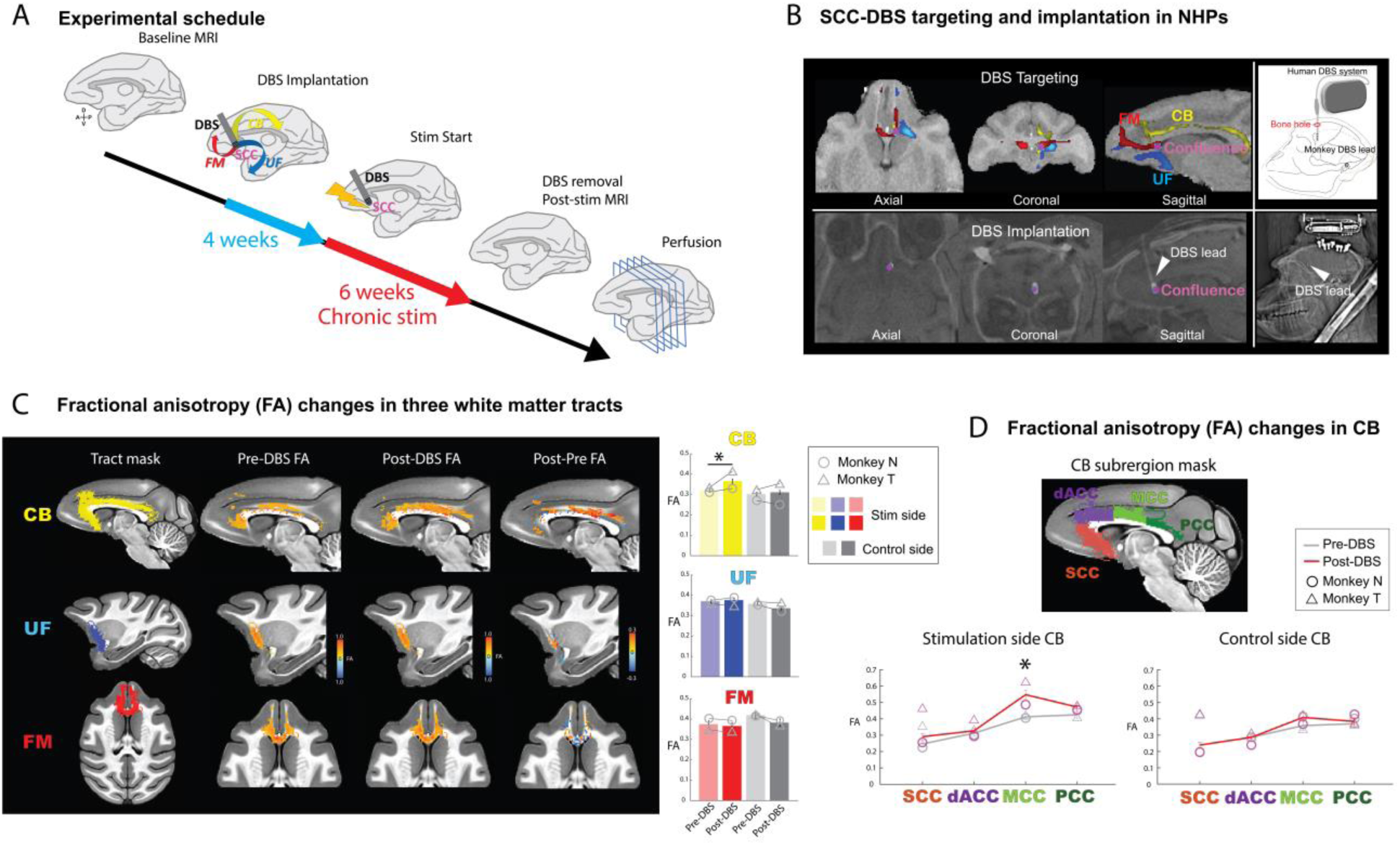
SCC-DBS changes fractional anisotrophy in the middle cinglum bundle. A) Experimental timeline. Baseline T1, diffusion weighted images and resting state functional MRI were collected for each subject. Using the baseline diffusion weighted images, the SCC-DBS target, i.e. the confluence of the cingulum bundle, the uncinate fasciculus and the forceps minor, was visualized. Using a stereotactic approach, a SCC-DBS lead was implanted to target the confluence in one hemisphere. After a four week recovery period, chronic stimulation for six weeks was delivered. Just after removal of SCC-DBS, diffusion and functional MRIs were collected. On the day after MRI, the animal’s brains were extracted and prepared for histological assessment. B) Identification of white matter tracts and confirmation of SCC-DBS lead location. Top-left: Visualization of the confluence visualization of CB, FM and UF made from overlapped images of the three reconstructed white matter tracts using probabilistic tractography on DWI (monkey T) and a T1w pre-DBS surgery structural image. Pink circle denotes the confluence of the three white matter tracts: cingulum bundle (yellow), uncinate fasciculus (blue), forceps minor (red). Top-right: The scheme of SCC-DBS implantation. A miniaturized custom-made DBS lead was implanted into the confluence through a small craniotomy and connected to a human DBS system. Bottom-left: Post-DBS surgery CT confirmation of DBS lead location. The post-DBS implantation CT image (monkey T) is overlaid on a pre-DBS implantation T1w image. Bottom-right: The CT scout view of SCC-DBS implanted macaque head. C) Fractional anisotropy (FA) in the three white matter tracts before and after DBS. Extracted FA values are shown from the cingulum bundle (CB, top), the uncinate fasciculus (UF, middle), and the forceps minor (FM, bottom). The second column from left shows pre-DBS FA value, the third column is post-DBS FA value, and the most right column is the subtraction image of post-DBS FA and pre-DBS FA values. The color indicates FA value. The bar graph shows mean (+/- SEM) FA value of pre and post-DBS in each tract in each hemisphere. Each bar represents the averaged FA value extracted from voxels included in each white matter tract mask. Colored bars indicate the mean FA value in the tract mask on the stimulated hemisphere (yellow for CB, blue for UF, red for FM) and the gray bars are those on the control hemisphere. Lighter color/gray bars represent pre-DBS FA and darker bars demonstrate post-DBS FA. D) FA value changes after SCC-DBS in subparts of the cingulum bundle. Mean (+/- SEM) FA value in four subparts (SCC, dACC, MCC and PCC portion) of the CB before and after stimulation in stimulated (bottom left) and control hemisphere (bottom right). Symbols represent individual subjects (circle/triangle, monkey N/T, respectively). * p < 0.05, Tukey-Kramer test. Error bars are SEM.

Fractional anisotropy (FA), derived from diffusion-weighted images (DWI), is a measure of white matter integrity (*35–37*) and has been shown to correlate with changes in white matter structure in humans and animals (*38*, *39*). We therefore obtained whole-brain DWI before SCC-DBS lead implantation and after six weeks of stimulation. We extracted FA values from the probabilistic reconstructions of the CB, uncinate fasciculus and forceps minor to quantify *in vivo* changes in white matter integrity following SCC-DBS. We then directly compared FA values within the tracts by subtracting the pre-surgery FA from post-stimulation FA values (**Fig. 1C**).

We found that there was a selective increase in FA in the cingulum bundle in the stimulated hemisphere, particularly the mid-cingulate portion, after six weeks of SCC-DBS (3-way repeated measures ANOVA, F_(1,3187)_ = 4.64, p = 0.0312, post-hoc tests on tracts Tukey-Kramer test, p = 0.022, **Fig. 1C** top row, **Fig. S2** left). No consistent changes were apparent in either the stimulated uncinate fasciculus or forceps minor (**Fig. 1C**, middle **and** bottom row **Fig. S2**, middle and right) or in the same tracts within the control hemisphere (**Fig 1C** bottom row and **Fig. S3 and S4**). Based on these findings, we next aimed to determine whether specific sub-regions of the CB showed major FA changes after SCC-DBS. Comparing the FA values before and after six weeks of DBS in four CB subregions revealed that only the midcingulate region on the stimulated side showed an increase in FA (3-way repeated mesures ANOVA, F_(1,1357)_ = 5.42, p = 0.020, post-hoc analyses, Tukey-Kramer test, p = 0.0017, **Fig. 1D** and **Fig. S4**).

In summary, *in vivo* neuroimaging indicate that SCC-DBS modified the white matter integrity of the CB after six weeks of stimulation. This anatomical change was specific to the midcingulate portion of the CB that is spatially remote from the site of stimulation delivery. Thus, SCC-DBS is associated with a selective macro-level remodeling of white matter distant from the stimulation site in a tract that connects the SCC with dorsal and posterior parts of the cingulate cortex (*40–42*).

### SCC-DBS induces increased myelinating oligodendrocytes and the degree of myelination

Prior work has shown that FA is associated with underlying white matter microstructure, however, such macro-level estimates are unable to reveal what specifically underlies this measure (*43–45*). Thus, we next sought to establish whether, and how, the SCC-DBS induced FA change was related to cellular-level changes in white matter in the midcingulate cortex. First, we investigated how stimulation influenced the proliferation of oligodendrocytes, especially myelinating oligodendrocytes in the midcingulate portion of cingulum bundle where FA increases after SCC-DBS were found, by conducting immunolabeling against CC-1, an antibody that preferentially labels myelinating oligodendrocytes (**Fig. 2A**). The proportion of CC-1 positive cells in the whole DAPI stained population was higher in the stimulated CB compared to the control side (*CC1/DAPI ratio:* one-way repeated measures ANOVA, F_(1,108)_ = 9, p = 0.0034, **Fig. 2B**, **Fig. 2C**, **Fig. S5**). Such a change was not observed in the portion of the CB adjacent to dorsal anterior cingulate, matching the lack of change in FA (**Fig. S6**). Thus, SCC-DBS was associated with an increase in the proliferation of oligodendrocytes specifically in the mid-cingulum bundle at the same location where we found a change in FA, and distant to where stimulation was delivered.

**Figure 2:**
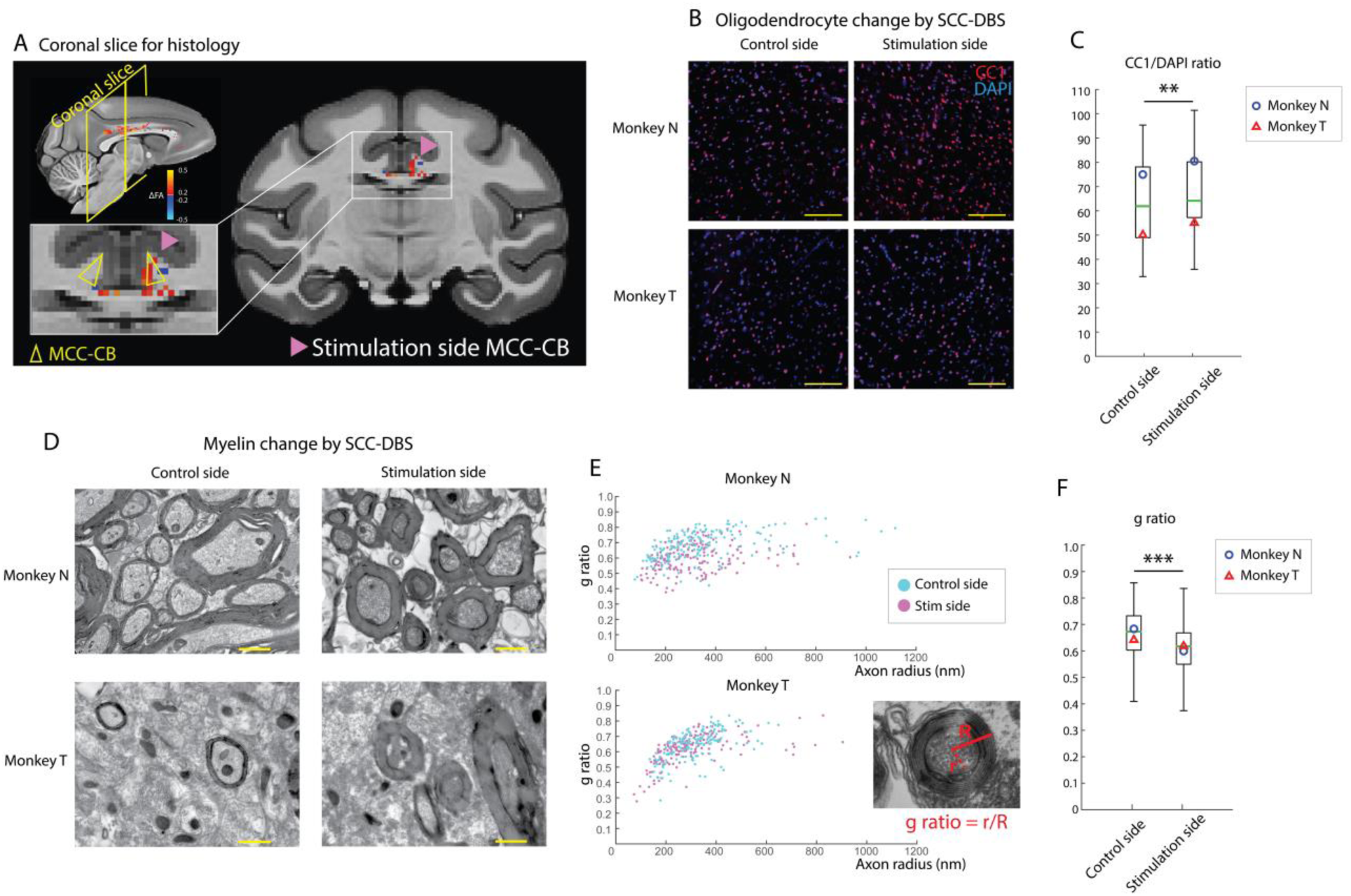
Cellular-level remodeling of in the mid-cingulum bundle by SCC-DBS. A) Sagittal and coronal images showing where changes in cellular level markers of white matter structures were assessed. B) Double stained images with CC1 (marker for myelinated oligodendrocytes, red) and DAPI (marker for nuclei, blue, x20 magnification) from monkeys N and T (top and bottom rows, respectively) for control (left) and stimulated hemispheres (right). Yellow scale bars indicate 100μm. C) Box plot of the ratio of CC1 and DAPI positive cells (%). Green line is the median of the CC1/DAPI ratio. Symbols denote means for each subject (circle/triangle, monkey N/T, respectively). **p < 0.01. D) Electron microscopy images of myelinated axons in control (left) and stimulated hemisphere (right) from the midcingulate portion of CB in monkey T. Yellow scale bars indicate 800nm. E/F) Scatter and box plot of g-ratio measurements from control (blue) and stimulated hemispheres (red). *** p < 0.001 from one-way repeated measures ANOVA.

Next, we aimed to establish whether SCC-DBS also changed the morphology of myelination within this region. Alterations in the thickness of the myelin sheath around an axon impact the conduction of signals along that axon, as well as neural synchrony (*46–50*). If stimulation is increasing FA in the mid-cingulum bundle, this should be associated with increased myelin thickness (*51*). Using electron microscopy, we measured the ratio between the inner and outer diameter of a random selection of myelinated axons within the mid-cingulate (**Fig. 2D**). This ratio, known as the g-ratio (**Fig. 2E** right bottom), measures the degree of myelination for each axon, allowing for the degree of myelination within a white matter tract to be quantified. In the stimulated hemisphere, the g-ratio in the mid-cingulum bundle was decreased compared to the control hemisphere (one-way repeated measures ANOVA, F_(1,696)_ = 58.98, p < 0.001, **Fig. 2D-F**), meaning that the degree of myelination was increased compared to the unstimulated control hemisphere.

Taken together these findings provide histological confirmation that SCC-DBS enhances myelin remodeling in the midcingulate portion of cingulum bundle and provides a direct biological basis for the increased FA seen after six weeks of SCC-DBS.

### Whole brain functional connectivity changes following SCC-DBS

Our macro and micro-level structural analyses of white matter indicate that the cingulum bundle was remodeled by SCC-DBS. The degree of myelination within a white matter tract is known to affect the conduction velocity of signals in that tract as well as the level of neural synchrony between areas that connect through that tract (*48*, *52*). Thus, based on the relationship between structure and function, we would expect that SCC-DBS should be associated with changes in fMRI functional connectivity (FC) between the stimulated SCC and regions in the networks that receive anatomical connections via the CB. We analyzed how SCC-DBS altered resting-state functional MRI (rs-fMRI) using a seed-based ROI analysis comparing FC between the stimulated SCC and the rest of the brain. Prior to stimulation, SCC showed positive connectivity to the ventromedial frontal cortex (vmFC), the dorsal anterior cingulate cortex (dACC), the posterior cingulate cortex (PCC), superior temporal gyrus (STG), hippocampus (HIP), amygdala (AMY) and dorsolateral prefrontal cortex (DLPFC). By contrast, SCC demonstrated negative connectivity to insula and somatosensory cortex before stimulation (**Fig. 3A**, top, **Fig. S7**, top, **Fig. S8**).

**Figure 3:**
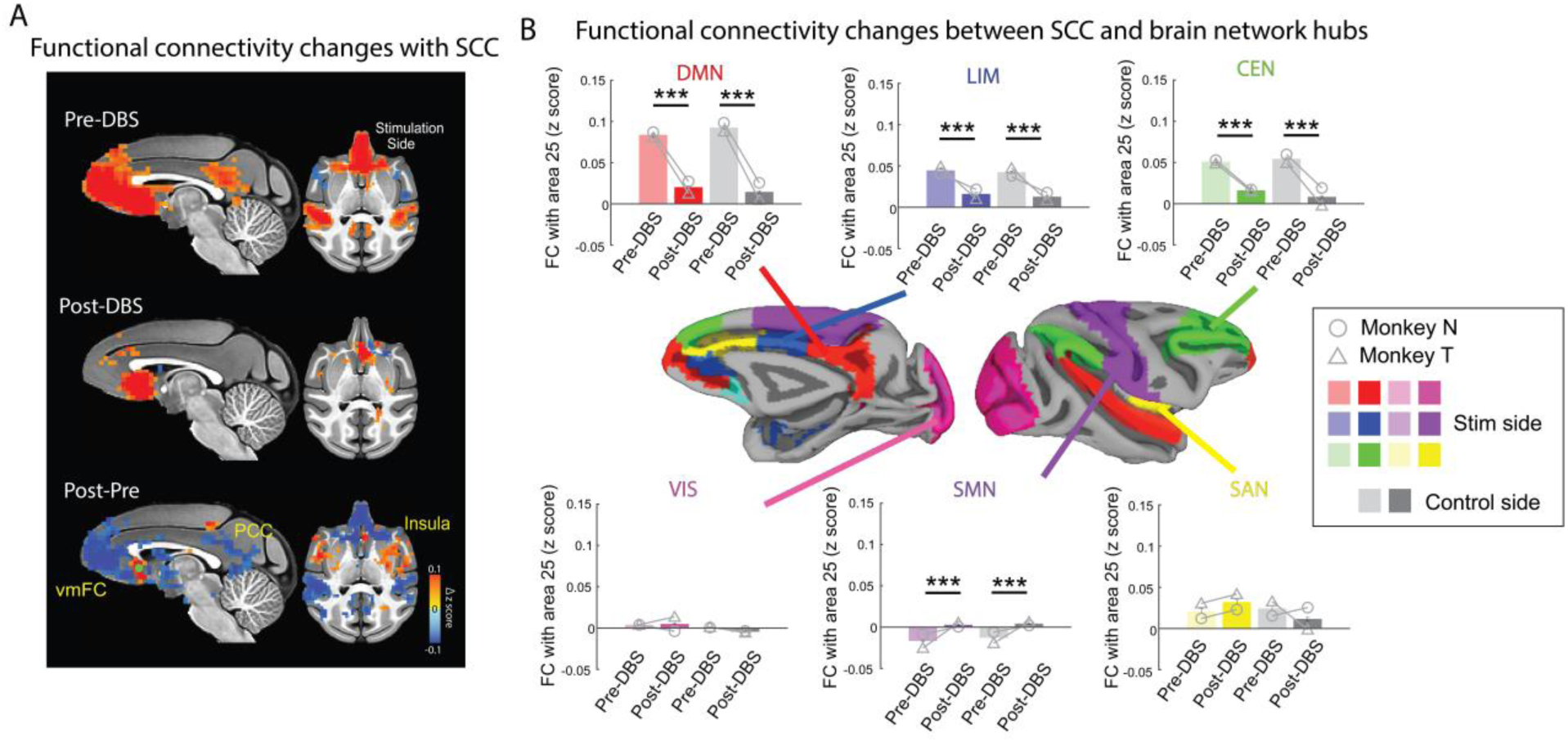
SCC-DBS associated changes in functional connectivity. A) The mean whole brain functional connectivity (FC) with area 25 shown on sagittal (left) and axial (right) images before (top row) and after (bottom row) six weeks of SCC-DBS stimulation, as well as and the difference between post and pre (bottom row). Sagittal images of FC from the stimulated hemisphere are shown. Images are thresholded for FC with p < 0.05 and cluster size >20. The green circle represents the location of the active contact of SCC-DBS lead. B) Intrinsic brain network-level changes in FC after six weeks of SCC-DBS. Changes in FC between the stimulated area 25 (purple region) and six brain network hubs (hubs of the default mode network (DMN), the limbic network (LIM), the salience network (SAN), the central executive network (CEN), the sensorimotor network (SMN) and the visual network (VIS)) are shown. Symbols on the plots represent individual monkey mean normalized FC values (circle/triangle, monkey N/T, respectively). Error bars show SEM. * p < 0.05, ** p < 0.01, *** p < 0.001. Tukey-Kramer test.

Following six weeks of SCC-DBS there was a marked reduction in functional connectivity of the SCC with the posterior cingulate cortex and ventromedial frontal cortex in addition to hippocampus, amygdala and dorsolateral prefrontal cortex (**Fig. 3A**, bottom, **Fig. S7**, bottom, **Fig. S8**, **Fig. S9**, 3-way repeated measures ANOVA, F_(1,26036)_ = 185.66, p < 0.001, post hoc Tukey-Kramer tests: vmFC, PCC, STG, HIP, AMY, DLPFC: p < 0.001, dACC: p = 0.0015). Most of these brain regions receive projections via the CB (*40*, *42*), and exhibited a positive correlation before stimulation. However, functional changes were not limited to these CB projection-related areas or decreases in connectivity; insula, somatosensory cortex and superior temporal gyrus all exhibited increases in FC with the SCC (**Fig. 3A**, post hoc Tukey-Kramer tests: anterior-dorsal insula, somatosensory cortex: p < 0.001). Despite unilateral DBS and unilateral localized white matter remodeling (**Fig. 1**), functional connectivity changes were also apparent in the unstimulated control hemisphere (**Fig. 3B**, **Fig. S8 and S9**). This matches findings that functional connectivity is not solely defined by structural connectivity (*53*) and that unilateral SCC-DBS, in humans, bilaterally impacts neural activity (*54*, *55*).

To investigate network-level functional connectivity changes, we used a hub-based approach (see **Materials and Methods**), calculating FC between SCC and each of six intrinsic networks: the default mode network, the salience network, the limbic network, the central executive network, the sensorimotor network and the visual network. After chronic stimulation, the default, limbic, and central executive networks showed significantly decreased FC with SCC (post hoc Tukey-Kramer tests: the default mode network: p < 0.001, the limbic network: p < 0.001, the central executive network: p < 0.001) (**Fig. 3B**). In comparison, there was no change in relation to the visual network or areas within the visual network (**Fig 3B** and **Fig. S9**). Such a pattern of changes fits with the idea that SCC-DBS is influencing activity in areas that connect via the CB which includes the default mode, central executive, and limbic networks (*56*, *57*).

Interestingly, networks not directly connected through the CB also exhibited functional changes, such as the sensorimotor network which showed an increase in connectivity with SCC (post hoc Tukey-Kramer tests: the sensorimotor network: p < 0.001) (**Fig 3B**). Meanwhile, the salience network did not exhibit consistent changes (**Fig. 3B**, yellow network), with the insula cortex showing an increase, and the dorsal anterior cingulate cortex showing a decrease in connectivity with SCC (post hoc Tukey-Kramer tests: anterior-dorsasl insula: p < 0.001, dACC: p = 0.0015) (**Fig. 3B** and **Fig. S9**). This highlights that SCC-DBS did not have a uniform effect on all nodes within an intrinsic network and appears to alter the balance of connectivity within the salience network.

Taken together, SCC-DBS was associated with a selective shift of functional connectivity between SCC and the rest of the brain, including areas that are not directly anatomically linked to SCC through the cingulum bundle. This pattern of effects supports prior neuroimaging investigations of depression that implicate altered connectivity between SCC and the default and salience networks (*30*, *58–65*).

## Discussion

Our results indicate that SCC-DBS is associated with focal white matter structural remodeling in the cingulum bundle as well as a selective shift in the connectivity of multiple functional networks. Further, our findings reveal the potential importance of remodeling of the cingulum bundle for effective DBS treatment for depression as well as the potential importance of white matter plasticity for DBS therapies in general.

Prior work has shown that white matter remodeling can be induced by electrical, optogenetic or chemogenomic stimulation of neurons in gray matter (*66–68*). Our data reveal that DBS targeted to white matter using clinically effective parameters in primates induces both macro and micro remodeling of white matter. Our findings also complement prior studies that have pointed to a role for dysfunction in white matter as contributing to the pathophysiology of depression (*69–73*). Notably, abnormal white matter integrity in the the cingulum bundle was recently found to be positively correlated with symptom severity in patients with treatment resistant depression (*32*). Additionally, treatment of depression with ketamine is associated with increased FA in the cingulum bundle (*74*, *75*). These data point to the long-term therapeutic effects of SCC-DBS being mediated at least in part by its effects on white matter. These data could also indicate that treatment resistant depression is associated with abnormalities in the cingulum bundle and that promoting remodeling of specific white matter tracts contributes to DBS-mediated recovery.

Another question is how SCC-DBS induced changes in network function and anatomy dynamically interact. Based on the progressive response of patients to SCC-DBS (*3*, *24*, *28*), it is likely that this is a multi-step process. Brain networks are known to dynamically re-balance in order to optimize input and outputs between brain areas (*76–79*) and SCC-DBS causes rapid changes in brain activity especially in the default mode network (*30–33*, *80*). On the other hand, remodeling the white matter structures takes weeks (*81*, *82*), with its effects on network function likely being more sustained (*30*, *32*). As such, SCC-DBS may be associated with a rapid shift in the balance of functional brain networks followed by more protracted white matter structural remodeling to strengthen modified functional connections or potentially to compensate for them.

Why SCC-DBS induced white matter remodeling was focused into the mid-cingulate portion of the cingulum bundle is not immediately clear. One possibility is that the change in white matter may be related to the alterations in functional brain networks. The mid-cingulate cortex has been characterized as a central hub for functional brain networks (*83*). Indeed, this area is where the default mode and salience networks overlap (*84*) and abuts the sensorimotor network. Thus, the confluence of these networks in the mid-cingulate and their DBS-induced shift in functional connectivity could be a key factor in driving white matter structural remodeling. Understanding the dynamic interplay between functional and anatomical mechanisms has the potential to further the development of SCC-DBS as well as identify biomarkers for personalized treatments for depression and other psychiatric disorders.

## Materials and Methods

### Subjects

Two male adult rhesus macaques (*Macaca mulatta*, monkeys N and T) 7-9 years of age were used. Subjects had ad libitum access to food and water and were maintained on a 12-hour light/dark cycle. SCC-DBS implantation, magnetic resonance imaging (MRI) scans (T1 weighted image (T1w), resting-state functional MRI (rsfMRI), and diffusion weighted image [DWI]), and histological experiments (immunohistology staining and electron microscopy analysis) were conducted in both subjects. All procedures were approved by the Institutional Animal Care and Use Committee of Icahn School of Medicine at Mount Sinai.

### Acquisition and processing of neuroimaging data

A pair of MRI sessions were conducted in each subject to acquire T1w, DWI, and rsfMRI, before SCC-DBS implantation as baseline scans (pre-DBS MRI) and immediately after the six week chronic stimulation (post-DBS MRI) (**Fig 1A**). Approximately 20 minutes prior to the post-DBS MRI, the DBS system, including the implanted DBS lead, was removed to avoid dropout and artifacts in the MRI images, as well as histological damage caused by the heating of the DBS lead and extension.

All scans were acquired on a 3 Tesla Siemens Skyra scanner (Siemens Healthineers, Malvern, PA) and were conducted under anesthesia. This approach was taken to reduce movement artifacts that could distort the neuroimaging results. Animals were first sedated using ketamine (5 mg/kg) and dexmedetomidine (0.0125 mg/kg), were then intubated, and then maintained on isoflurane (0.7 – 3% to effect). Throughout the acquisition of the scans, the plane of anesthesia was continuously monitored by certified veterinarian staff monitoring a suite of vital signs (pulse, SpO2, end-tidal CO2, capnograph, blood pressure, body temperature).

Using a human 32ch head coil (Siemens Healthineers, Malvern, PA), DWI (b=1000 s/mm^2^, TR 6900ms, TE 96.0ms, 72 direction, resolution 1.5mm isotropic voxel, 8 unweighted volumes per scan, multi-band acceleration =3, two opposite phase encoding = AP/PA) were acquired with the animal in the supine position. We collected six total runs of the DWI data, 3 with AP phase encoding and 3 with reversed phase encoding (PA). Two to three T1w images (0.5 mm isotropic, TR/TE 2500/2.81 ms, flip angle 8°) were also collected using the same position and coil system. For both DWI and structural scans the level of isoflurane was kept around 1-1.5% to ensure the stability due to the use of a stronger gradient system typical for sequences in DWI.

Resting-state functional MRI (rsfMRI) scans were performed using the same protocol we have previously reported (*85*), using a custom-built 4-channel phased array transmit/receive coil (Windmiller-Kolster Scientific, Fresno, CA). Unlike the structural and DWI scans, anesthesia was maintained on a low level of isoflurane (0.7-0.9%). The use of low-level anesthesia allows for the preservation of resting-state networks (*85–89*). To reduce stimulation, the animal’s head was not restrained but supported with towels and flexible bandages to allow for stability with minimal discomfort. To minimize the effect of physiological changes in the neural activity, vital signs (pulse, SpO2, end-tidal CO2, capnograph, blood pressure, body temperature) were continuously monitored and maintained throughout an experimental session. A session-specific 3D T1w image (0.5 mm isotropic, TR/TE 2500/2.81 ms, flip angle 8°) was acquired. Following injection of monocrystalline iron oxide nanoparticle (MION, i.v.), to improve the contrast-to-noise ratio of the functional data (*90*, *91*), six functional scans were obtained (Echo Planar Images (EPI): 1.6 mm isotropic, TR/TE 2120/16ms, flip angle 45°, 300 volumes per each run). At the completion of all scans, subjects were extubated and continuously monitored until they were able to sit unaided.

### DWI data preprocessing

A DWI data preprocessing pipeline was established by modifying standard human DWI analysis pipelines (*22*) using AFNI(*92*), FMRIB Software Library (FSL) (*93–95*), Advanced Normalization Tools (ANTs) (*96–99*), ITK-SNAP (*99*), and MRtrix3(*100–104*)(*95*). Raw DWI data was first converted into the MIF file format. Then, it was denoised based on principal component analysis in order to improve the signal noise ratio. After this, the data from the two different phase encoding directions (AP and PA) were merged, and then susceptibility-induced distortions, eddy currents and the bias b1 field were corrected. Non-diffusion-weighted images were extracted, averaged, and converted into NIFTI file format. After brain extraction, images were aligned with session specific average T1w linearly. Then, diffusion tensors were fitted to each voxel using DTIFIT (*105*). Voxel-wise crossing-fiber model fitting of diffusion orientations was performed using FSL’s BedpostX (*106*).

### Identification of white matter tracts for DBS targeting

In the current study, we chose to target DBS to the confluence of three white matter tracts near to the SCC. We chose this location based on data showing that targeting this point in the brain is associated with positive clinical outcomes for treatment resistant depression (*22*, *23*). To do this we reconstructed the cingulum bundle, uncinate fasciculus, and forceps minor separately using probabilistic tractography (FSL’s ProbtrackX) based on our pre-DBS DWI (*106*). White matter tract reconstruction was conducted in each individual’s space, therefore removing issues of warping to a common space. All seed masks, waypoint masks and exclusion masks were made with session specific principal eigenvectors. These masks were created based on previous anatomical studies, tract-tracing studies, and diffusion tractography studies (*40*, *41*, *107–112*). Masks were defined as follows:

Cingulum bundle (CB) seed mask: The CB seed mask was drawn in a coronal plane at the level of the genu of the corpus callosum by capturing the white matter dorsal and medial to the corpus callosum. The seed mask showing the curve connecting subcallosal anterior cortex to parahippocampal region was selected (*42*). Additionally, for monkey N, the seed mask for the subcallosal part of the CB was drawn separately in a sagittal plane to visualize the subcallosal part of CB more clearly.

Uncinate fasciculus (UF) seed mask: UF seed mask was made on axial slices in the white matter of the anterior temporal lobe. The final seed mask was chosen to account for the strong curve of the fibers from dorsal-ventral to anterior-posterior orientation as they enter the frontal lobe (*107*).

Forceps Minor (FM) seed mask: The seed mask for FM was drawn in one hemisphere’s frontal lobe in the coronal plane near the frontal pole, with the target mask placed in the contralateral hemisphere in the same coronal plane (*111*).

We used a coronal waypoint mask at the level of the rostral genus of the corpus callosum to ensure that the fibers emanating from each seed projected to the frontal cortex for the CB and UF (*107*). For the FM, we used a waypoint mask draw on the posterior part of the FM. To reduce false positive artifacts, we used a set of exclusion masks that included (1) a basal ganglia mask and a seed mask for the internal capsule to avoid leaking into the anterior limb of the internal capsule, (2) a posterior mask for the UF and FM, to avoid false positive tracts running in a posterior as opposed to an anterior direction (3) an interhemispheric mask separating the brain into left and right hemispheres, except for the FM, (4) a separate frontal lobe interhemispheric mask for the FM, which was the anterior part of the interhemispheric mask covering only the frontal lobe anterior to the corpus callosum, and (5) the other tract seeds allowing the CB, UF and FM to be exclusive (for example, the seeds for UF, FM, and the internal capsule were used as exclusion masks for the CB).

Probabilistic tractography was run in each monkey’s diffusion space, using the above seeds, a waypoint mask, and exclusion masks (maximum steps per sample = 3200; samples = 10,000; step size = 0.1 mm; curvature threshold = 0.2). Each tract was log-transformed to account for the exponential decrease of visitation probability with distance and normalized by dividing each voxel’s value by the maximum value across the tract, removing the potential bias of differences in numbers of streamlines between tracts. Normalized tracts were subsequently thresholded with minimum and maximum values equal to 0.7 and 1 respectively matching previous methods (*107*).

The confluence of three tracts was determined in each animal as the location where the separately calculated tracts overlapped in an 8-voxel square (3×3×3mm). To confirm whether the confluence seed was the optimal SCC-DBS target, we compared the tracts going through the confluence with nearby seeds of the same size. This was done using probabilistic tractography without the exclusion or waypoint masks since it clarifies the possible tracts which DBS can have effects. We decided the final target (3×3×3mm) as the seed that showed (1) ipsilateral CB, (2) ipsilateral UF, and (3) bilateral FM including the posterior part that connects left and right area 25 and the anterior parts of the left and right frontal poles.

To counterbalance for the side of stimulation and also to ensure that there was clear overlap in the tracts at the point of stimulation, the confluence in the left hemisphere was selected in monkey N and the right-side in monkey T.

### SCC-DBS implantation surgery

A pre-operative MRI scan (T1w and DWI) was used in each animal to target the confluence of three white matter tracts (cingulum bundle, uncinate fasciculus and forceps minor, see *DBS targeting*). Alignment between scan and surgical space was ensured using toothmarker registration of the subjects’ head relative to the stereotactic frame (*113*). As described above, the optimal DBS target was determined in the pre-DBS DWI space and aligned to the pre-DBS T1w space. The DBS lead was implanted in one hemisphere per subject (left in monkey N; right in monkey T). The other, unimplanted hemisphere was preserved as a non-stimulation control.

In a dedicated operating suite using aseptic procedures, anesthesia was induced using ketamine (5 m/kg) and dexmedetomidine (0.0125 mg/kg) and then maintained by isoflurane (2-3%). The skin, fascia and muscles were opened and retracted. A small craniotomy was made in the skull at the location above the confluence of the three tracts and an opening in the dura was made. A 22G stylet needle covered by 18G sheath needle was inserted into the brain to a point just above the target to make an insertion tract. The 22G stylet was then removed, and the custom-made mini-DBS lead (4 electrode connector type mini lead, NuMed Inc., Hopkinton, NY, USA) was inserted using stereotactic control through the 18G sheath into the insertion tract and placed at the location of the confluence. The sheath was then removed. The DBS lead was fixed in by filling the craniotomy with adhesive silicone (KWIK-SIL, low toxicity silicone adhesive, World Precision Instruments, Sarasota, FL, USA). For safety, the DBS generator (Activa SC for Monkey N, Percept PC for monkey T, Medtronic, Minneapolis, MN, USA) was set in a 3D printed case (Form2 (3D printer), Formlabs (3D designing software), Dental LT clear VC Biocompatible Resin (3D printing material), Formlabs Inc., Someville, MA, USA) that was fixed on the skull using acrylic cement (Lang Dental, IL, USA). The mini-DBS lead and DBS generator were connected by a human DBS extension (37086, Medtronic). The DBS generator was grounded in monkey N to a titanium screw (Veterinary orthopedic implants, Augustine, USA) implanted on the control side over the parietal cortex connected by a silver wire (0.004” diameter, Surepure Chemicals, Florham Park, NJ, USA). In monkey T, a second small craniotomy was made in the control side parietal skull and a second DBS lead was inserted in the epidural space as a grounding wire connected to the generator. The 3D printed case containing the DBS generator was filled with conductive gel (New Bio Technology Ltd., Israel). The muscles, fascia, and skin were then sutured closed.

The lead placement was confirmed by high-resolution CT (0.5mm isotropic CT, Force CT, Siemens Healthineers, Malvern, PA) two weeks after SCC-DBS implantation surgery. CT images were then aligned to the pre-DBS T1w scans for each subject. The CT defined location of the inserted DBS lead was then compared with our predefined optimal DBS target location (**Fig. 1C**).

### SCC-DBS chronic stimulation

Based on the CT scan, the closest contact to the optimal SCC-DBS target out of the four lead contacts was selected as the stimulation contact. After a 4-week postsurgical recovery period, we started 6 weeks of chronic stimulation (130Hz, 90μzsec, 5mA). We chose this stimulation regime as it has been shown to be effective parameters for human depression (*3*, *22*, *28*).

### Fractional anisotropy analysis

Fractional anisotropy (FA) was calculated from pre-DBS and post-DBS scans in each animal’s own space and aligned to the subjects’ T1-weighted structural scans (ANTs) (*99*). The warp parameter for registering the post-DBS T1 image to pre-DBS T1 image was calculated using the C3D nonlinear registration function, and then warps were applied to the post-DBS FA maps to align all the data to the same space. This method was used in order not to lose FA signal for each tract. Whole brain FA subtraction was conducted in pre-DBS T1 space for each monkey. The FA value was extracted from each voxel (1.5mm isotropic) in the white matter. Pre-DBS FA and post-DBS FA were compared for each tract (CB, UF and FM) using a 3-way repeated measures ANOVA (DBS (pre-/post-) × side (stim/control) × tract (CB/UF/FM)), followed by a post hoc Tukey-Kramer test, revealing the white matter tracts that were affected by SCC-DBS the most (Matlab, R2023a, Mathworks, Natick, MA, USA). Monkeys were modeled as a random effect.

To isolate where within the CB SCC-DBS caused changes in FA, we divided the CB mask into 4 subregions based on an anatomical atlas (D99 atlas; Saleem et al., 2021); SCC mask (CB adjacent to area 25 and area 32), dACC mask (CB adjacent to area 24), MCC mask (CB adjacent to area 24’) and PCC mask (CB adjacent to area 23). Similar to the above analysis comparing CB, UF and FM, pre-DBS FA and post-DBS FA in each CB subregion were compared using 3-way repeated measures ANOVA (DBS × side × subregion (SCC part/dACC part/MCC part/PCC part)), followed by Tukey-Kramer post-hoc test. Monkeys were modeled as a random effect.

To visualize the detailed FA value changes in each white matter tract, FA values within a tract were averaged for each slice. The CB mask was divided into two parts: an SCC dorsal/ventral (axial slices) part and an anterior/posterior (coronal slices) part for the remainder, because of its C shape, using the same definition above. The UF mask was also divided into two parts, the temporal (axial slices) part and the frontal-insula (coronal slices) part because of its C shape. The FM mask was divided into left and right hemispheres.

### EPI data analysis

Functional imaging data were preprocessed with a custom AFNI/SUMA pipeline (*85*, *114*, *115*). Raw EPI images were initially converted into NIFTI data file format and then into BIDS format (*116*). The T1w images were spatially normalized, skull-stripped (*117*), then aligned to atlas space (NMT-2.0; (*118*, *119*)). The images were first slice time corrected, then the first three TRs of each EPI were removed, and motion correction was applied. The EPIs were aligned to the within session T1w image and then warped to the standard NMT space. EPIs were blurred with an FWHM of 3 mm, and then converted to percent signal change. Finally, the motion derivatives from each scan along with cerebrospinal fluid and white matter signal were regressed out of the data. The residuals from this analysis were then used to compute the functional connectivity analysis using seed-based analysis.

The functional connectivity (FC) of the stimulated hemisphere’s SCC, area 25 (D99 atlas; (*119*)), was calculated(*119*) with other anatomically defined brain regions, in NMT space. Regions of interest, defined from the D99 atlas (*119*), CHARM atlas (*115*) and SARM atlas (*120*) were used. Target ROIs were: area 25 on the unstimulated side, area 32, dorsal ACC (dACC, area 24abc), middle cingulate cortex (MCC, area 24a’b’c’), posterior cingulate cortex (PCC, area 23, area 31), dorsal insula, hippocampus (HIP), amygdala (AMY), dorsolateral prefrontal cortex (DLPFC, area 8, area 9, area 46), ventromedial prefrontal cortex (vmFC, area 10mr, area 10o, area 10mc), superior temporal gyrus (STG), posterior parietal cortex (PPC, inferior parietal part of area 7), primary motor cortex (area 4), primary sensory cortex (area 1, 2, 3ab), supplementary motor area (SMA, area 6), primary visual cortex (V1), middle temporal lobe (MT, another visual network hub) (**Fig. S9**).

We calculated the Fisher’s z-transformed Pearson’s correlation between the average time series of the seed ROI and the time series of every voxel in the whole brain (AFNI’s 3dTcorr1D). Since pre-DBS EPI and post-DBS EPI are different sessions, the z-scores were normalized across session by dividing each voxels z-score the maximum z-score for each session. To statistically evaluate the effect of SCC-DBS on stimulated area 25 the normalized z-scores for pre-DBS and post-DBS in each target ROI were submitted to 3-way repeated measures ANOVA (DBS × side × target ROI) followed by post-hoc analysis (Tukey-Krammer test). Monkeys were modeled as a random effect. The FC changes were calculated by subtracting pre-DBS z-scores from post-DBS z-scores. To examine the brain network effects of SCC-DBS, hub brain region target ROIs were combined for each network respectively; PCC vmFC and STG for the default mode network, anterior-dorsal insula and dACC for the salience network, DLPFC and PPC for the central executive network, area 32, dACC, MCC, PCC, HIP and AMY for the limbic network (dACC was overlapped with the salience network and the limbic network, PCC was double counted in the default mode network and the limbic network), primary motor cortex, primary sensory cortex and SMA for the sensorimotor network, and V1 and MT for the visual network. To assess the statistical effect of SCC-DBS on brain network hubs, 3-way (DBS × side × brain network hubs) repeated ANOVA followed by post-hoc Tukey-Kramer tests were conducted. Monkeys were modeled as a random effect. The statistical significance was determined when the p-value was under 0.05.

### Behavior analysis

Behavioral data were collected while subjects were alone in their home cage for ten minutes without human intervention. In order to investigate the extent of behavior, i.e. the amount of movement, the ten minute video was divided into sixty bins (10 seconds per bin) and binarized such that bins with movement (walking, climbing, foraging etc.) were recorded as 1, and bins without movement were recorded as 0. We did the same analysis for foraging behavior. These analyses were conducted manually. We compared the pre-DBS (before stimulation start, after four weeks recovery period from DBS implantation) and the post-DBS (after six weeks stimulation) to establish the effect of SCC-DBS on natural homecage behavior. The binarized data was assessed by one-way repeated measures ANOVA. Monkeys were modeled as a random effect. The statistical significance was determined when the p-value was under 0.05.

### Histological processing

Animals were deeply anesthetized with a sodium pentabarbitol solution and perfused transcardially with 0.1 M phosphate buffered saline (PBS) and 1% paraformaldehyde (PFA) solution followed by 0.1 M PBS and 4% PFA solution. Brains were removed and immersed in 4% PFA solution overnight. Following post-fixation, brains were transferred to a solution of 2% DMSO and 10% glycerol in 0.1 M PBS for 24-48 hrs, then transferred to a solution of 2% DMSO and 20% glycerol in 0.1 M PBS for at least 24 hours. Cryoprotected brains were flash frozen in −80℃ isopentane. After flash freezing, the brains were removed, dried, and stored at −80℃ until sectioning. Tissue was cut serially in 30 μm steps for immunofluorescence staining and 120 μm (monkey N) or 200 μm (monkey T) for electron microscopy analysis in coronal sections on a sliding microtome equipped with a freezing stage (Leica SM 2010R, Leica Biosystems, Deer Park, IL, USA). Slices for immunofluorescence staining were stored at −20°C in antifreeze solution (20% Glycerol, 30% Ethylene glycol, 50% PBS). Slices for electron microscopy analysis were stored at 4°C in fixation solution (2% PFA, 2.5% glutaraldehyde, 0.1M Sodium Cacodylate Buffer (SCB) pH 7.4, Electron Microscopy Sciences (EMS), Hatfield PA, USA).

### Immunofluorescence of myelinating oligodendrocytes

Free-floating immunohistochemistry was performed for four coronal serial sections taken from midcingulate level with 750μm spacing for monkey N and 790μm spacing for monkey T. After washing extensively with phosphate-buffered saline (PBS) (10min for each wash step, 3 times), brain sections were immersed in target retrieval solution at 60°C for 30 mins (Target retrieval solution, Citrate pH6.1, Cat#S1699, Agilent Dako, Santa Clara, CA, USA, adjusted to pH 6.0 before use) as an antigen retrieval step. Then, after washing twice with PBS, these sections were incubated for 24 h with blocking solution (5% normal goat serum (Cat #PI31873, Invitrogen, Waltham, MA, USA) and diluted with PBS solution containing 0.3% Triton X-100 (TX-100)) at 4°C. After the blocking step, sections were incubated for 48 hours at 4°C with anti-CC1 primary antibody (anti-APC[CC-1] mouse monoclonal antibody, 1:5000, Cat# ab16794, RRID:AB_443473, Abcam, Cambridge, UK), the marker for the myelinating oligodendrocytes, diluted in the blocking solution. Sections were then washed and incubated for 2 hours at room temperature with goat polyclonal secondary antibody (IgG (H+L) Highly Cross-Adsorbed Goat anti-Mouse, Alexa Fluor™ Plus 555, 1:500, Cat# A32727, RRID:AB_2633276, Thermo Fisher Scientific, Waltham, MA, USA) diluted with PBS solution containing 0.3% TX-100. After a washing step with PBS, sections were stained with DAPI (DAPI nucleic acid stain molecular Probes, Cat# D1306, Invitrogen, Waltham, MA, USA, 0.0625Μg/ml diluted with PBS) for 10 minutes at room temperature followed by washing step with PBS. Sections were then dehydrated in ethanol (50% ethanol for 5 minutes, then 70% ethanol for 5 minutes) and immersed with autofluorescence eliminator reagent (Autofluorescence eliminator reagent, Cat# 2160, Millipore Sigma, Burlington, MA, USA) for lipofuscin staining for 10 minutes at room temperature. After washing with ethanol (70% ethanol for 3 minutes, then 50% ethanol for 1 minutes), the sections were mounted onto glass slides, air dried, and then cover slipped with prolong gold mountant (Prolong gold mountant, CAT #P36930, Invitrogen, Waltham, MA, USA). Images of the tissue were acquired on a confocal microscope (Zeiss LSM780 confocal microscope, Zeiss, Oberkochen, Germany). Seven non-overlapped images at 20x magnification per cingulum bundle were acquired (28 images per hemisphere per animal, 7 images for each hemisphere × 4 coronal sections). Cell counting for CC-1 positive cells and DAPI positive cells was conducted using FIJI software (version 1.54, Schindelin et al., 2012) using an adjusted threshold for each antibody for each monkey. This approach produces results that are essentially identical to the manual counting. The same threshold was applied for each section. In addition to counting the number of CC1 positive cells and DAPI positive cells per field, the ratio of the number of CC1 positive cells divided by the number of DAPI positive cells (the ratio of CC1/DAPI) was calculated in order to normalize the number of oligodendrocytes to the total number of cells in the population of cells.

The number of CC1 positive cells, the number of DAPI positive cells, and the ratio of CC1/DAPI for the stimulation side CB and the control side CB were compared using one-way repeated measures ANOVA. Monkeys were modeled as a random effect. The statistical significance was determined when the p-value was under 0.05.

### Electron microscopy analysis of myelin thickness

After two to three months in fixation solution (2% PFA, 2.5% glutaraldehyde, 0.1M Sodium Cacodylate Buffer, described in *Histology preprocessing* section), the midcingulate portion of CB in the coronal sections, adjacent to the sections used for immunofluorescence staining, was blocked and prepared for electron microscopy. Taking these sections ensured that we were looking at parts of the CB where we had assessed CC-1 staining.

Tissue blocks were washed in sodium cacodylate buffer, and then placed in 0.1% tannic acid/SCB for 30 minutes. After washing with SCB, the tissue blocks were incubated with 2% osmium tetroxide and 1.5% potassium ferricyanide diluted in SCB for 30 minutes in a dark setting. Then tissue blocks were washed with filtered distilled water and incubated with 1% thiocarbohydrazide in distilled water for 30 minutes. After washing with distilled water, the tissue blocks were immersed in 1% uranyl acetate in distilled water for 1 hour in the dark followed by a washing step with distilled water. Lastly, tissue blocks were incubated in lead aspartate staining solution (0.0066g lead nitrate/ 1ml aspartic acid solution) for 30 minutes at 60℃ and washed with distilled water again.

Tissue blocks were dehydrated in ascending ethanol concentrations and washed with100% ethanol three times (10 to 15 minutes incubation for each dehydration step) at room temperature. Tissue blocks were rinsed in 100% ethanol mixed 1:1 with propylene oxide for 10 minutes, followed by 100% propylene oxide for 10 min. The tissue was then infiltrated with EMbed-812 resin (EMS) mixed 2:1 with propylene oxide for 1 hour, followed by 1:1 EMbed-812: propylene oxide for 1 hour and then 1:2 EMbed-812: propylene oxide for 1 hour at room temperature. The tissue blocks were next placed into EMbed-812 for 2 hours, then placed into flat molds in the coronal orientation, filled with fresh resin, and and placed into a 65°C oven for 72 hours. Ultra thin sections (80 nm) were taken on a Leica UltracutEM UCT (Leica, Wetzlar, Germany) were mounted on Formvar/carbon coated slot grids (EMS, FCF2010-Cu) or 100 mesh Cu grids (EMS, FCF100-Cu). To evaluate the myelin thickness, eight to ten nonoverlapping images were taken per hemisphere for each animal respectively at 4,000 × to allow for an analysis of the g-ratio. Four measurements were recorded for each myelinated axon: the longest axon diameter, the shortest axon diameter, the longest myelin width, and the shortest myelin width. To calculate the g-ratio, the average diameter for each axon was divided by the average axon diameter plus twice the average myelin width (Dupree, Polak, Hensley, Pelligrino, & Feinstein, 2015; Murcia-Belmonte et al., 2016). Myelin regions that exhibited fixation artifacts or noncompaction were excluded from the analysis. Some axons were sectioned obliquely and demonstrated that the ratio of the longest axon diameter to the shortest axon diameter was over two. These axons were excluded in order to focus on the CB fibers running in the anterior-posterior direction and connecting subcallosal anterior cingulate cortex and posterior cingulate cortex. On average, the g-ratios of 175 myelinated axons were analyzed per hemisphere for each animal. Measurements of g-ratio were confirmed by a second, blinded rater.

The g-ratio in the stimulation side CB and the g-ratio in the control side CB were compared using one-way repeated measures ANOVA. Monkeys were modeled as a random effect. The significance was defined when p value was under 0.05.

## Acknowledgments

We would like to thank Dr. Paula Croxson for providing the foundation on which this work was built, Dr. Jungho Cha, Haneul Song and Jack Gomberg for advising on MRI analysis, Dr. Akbar Alipour for support on MRI settings, and Niranjana Bienkowska for assistance with data acquisition. For support on fMRI data pre-processing and analysis we thank Drs Paul Taylor and Alex Franco, respectively. For help with immunohistology procedure establishment we thank Dr. Danielle Beckman and Dr. Giovanne B. Diniz. For assistance with immunohistology data analysis and electron microscopy analysis, we thank Liza London, and Anatoli Velikov. For support on 3D printing, we also thank for Dr. Feng-Kuei Chiang. For help with animal perfusion reagent preparation, we thank Allison Sowa. For instruction about the surgical procedure of DBS implantation, we thank Dr. Brian H. Koppel and Dr. Josue M. Avecillas-Chasin. Finally, we thank the veterinary and animal care staff at Mount Sinai for their expertise and support.

## Author contributions

SHF, BER, HSM and PHR designed the study. SHF, AF and PER conducted the DBS implantation surgeries. SHF collected and ananalyzed the behavioral data. SHF, AF, CE, GV, LF, KS and BER established the MRI acquisition protocol. SHF, DF, KS and BER made MRI preprocessing pipelines. SHF analyzed the imaging data. SHF, WGJ and PER extracted and processed the brains. SHF and EA conducted and analyzed immunohistological assessment. SHF and AS analyzed electronmicroscopy analysis. Funds were acquired by SHF, BER, HSM and PER. This project was supervised by BER, HSM and PER. SHF, BER and PHR wrote the original draft. All authors edited the paper.

## Supplementary Figures

**Supplemental Figure 1:**
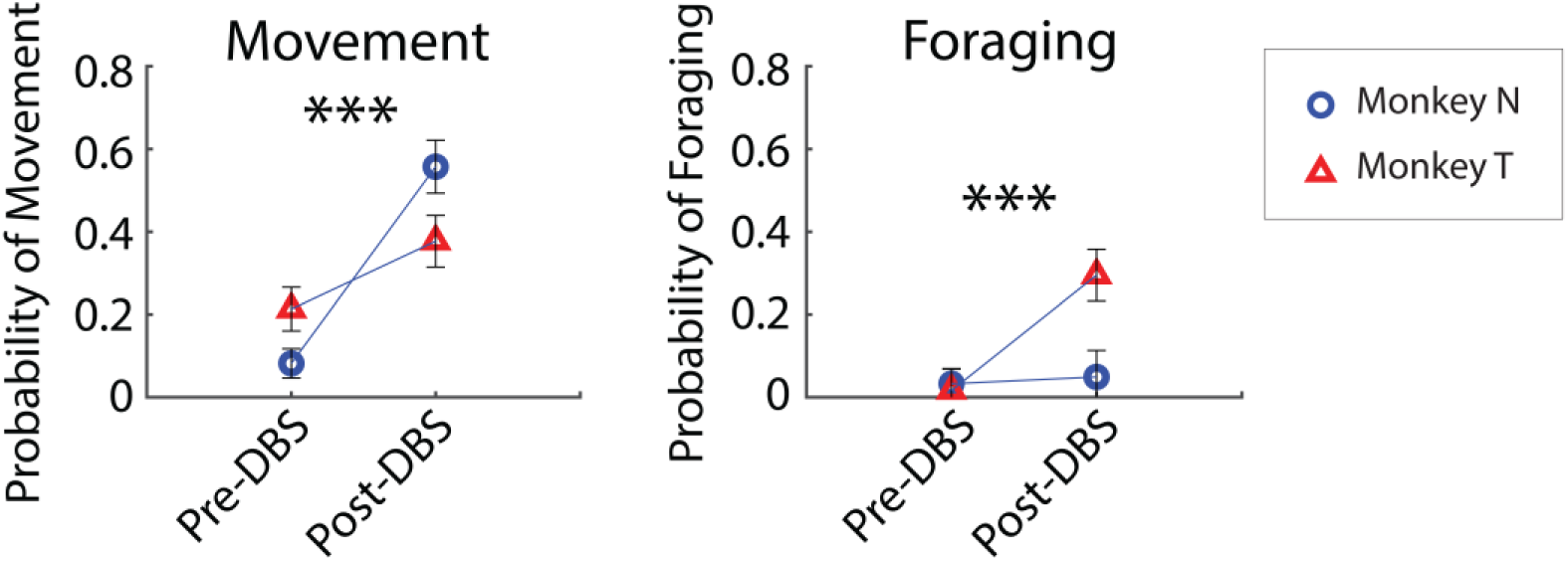
Naturalistic behavioral changes after SCC-DBS stimulation. The probability of movement (left) or foraging (right) observed in a 10-second bin among 60 bins (10 minutes) during either the pre or post SCC-DBS stimulation sessions. Symbols indicate probability of observed behavior for each animal (blue circle, monkey N, red triangle, monkey T). Error bars show SEM. A one-way repeated measures ANOVA was conducted to compare the effects of SCC-DBS stimulation. We observed an increase in both behaviors as a result of SCC-DBS stimulation. *** p < 0.001.

**Supplementary Figure 2:**
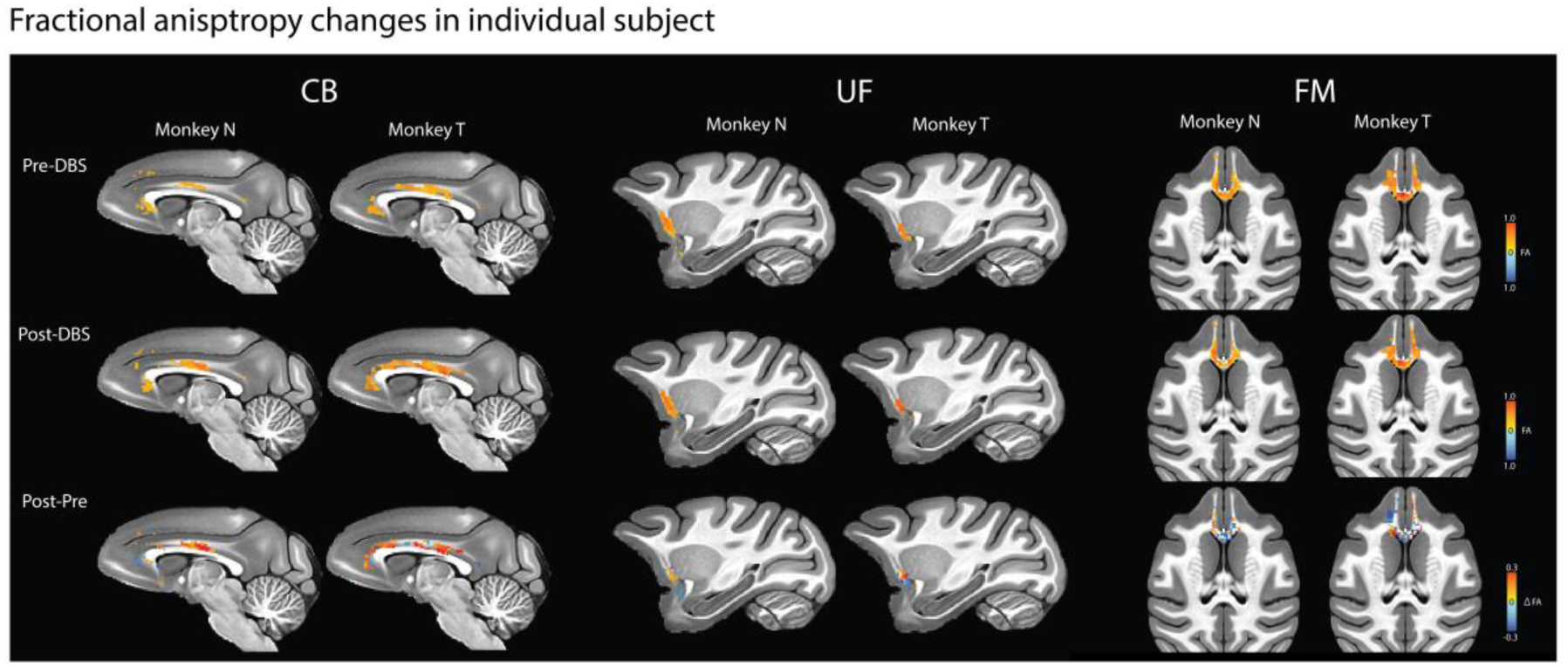
Fractional anisotropy changes in individual subjects. FA value in the cingulum bundle (CB, left), uncinate fascicle (UF, middle), and forceps minor (FM, right) in each subject. The extracted FA value in each white matter tract of interest is shown in NMT space. The top row is pre-DBS FA value, the middle row is post-DBS FA value and the third row is the subtraction image of FA value (post-DBS FA value - pre-DBS FA value) in each panel. The color indicates FA value.

**Supplementary Figure 3:**
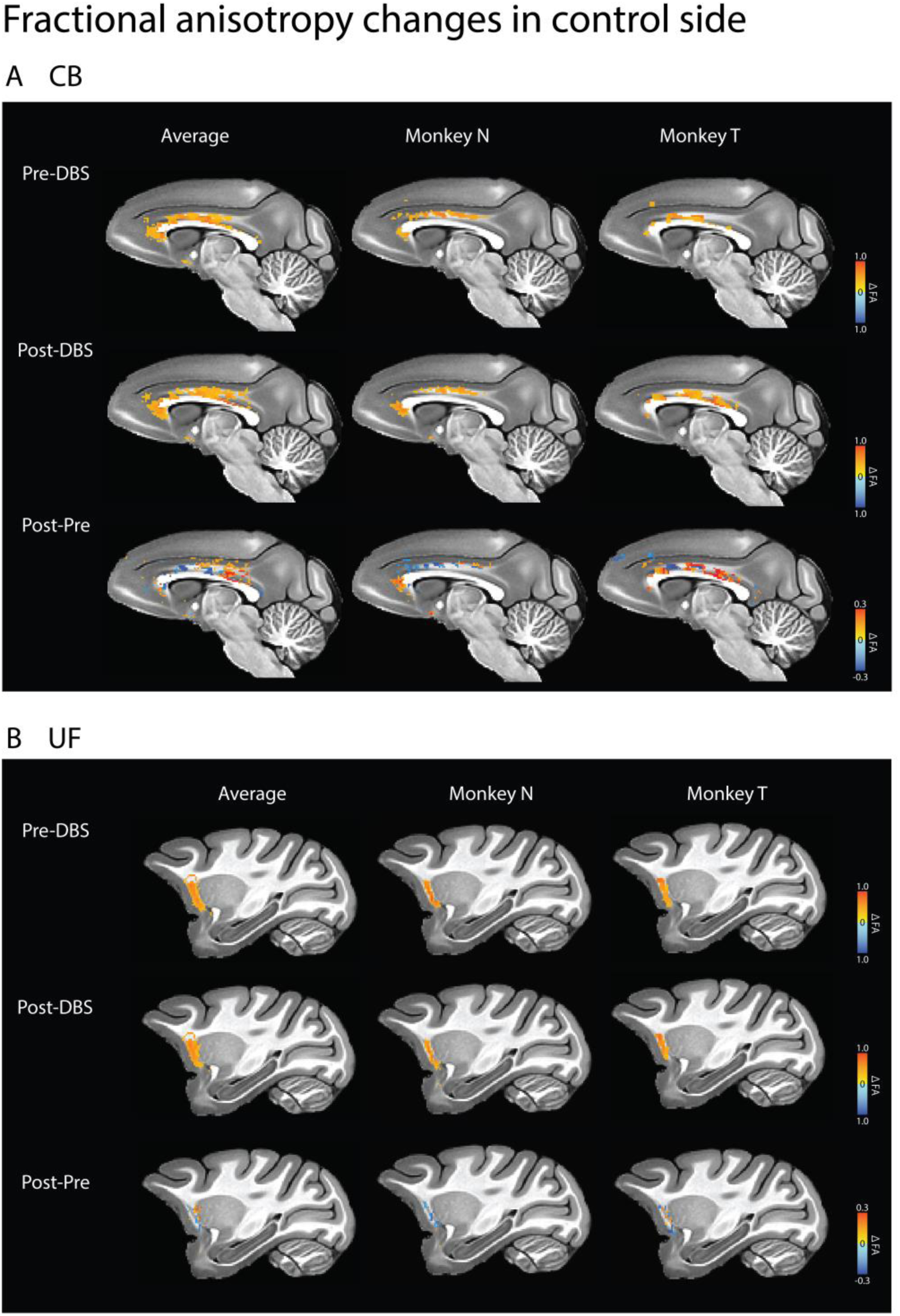
Fractional anisotropy in unstimulated control hemisphere. FA value in the CB (A, top) and UF (B, bottom) in the control hemisphere. The extracted FA value in each white matter tract of interest is shown in NMT space. The top row is pre-DBS FA value, the middle row is post-DBS FA value and the third row is the subtraction image of FA value (post-DBS FA value - pre-DBS FA value) in each panel. The color indicates FA value.

**Supplementary Figure 4:**
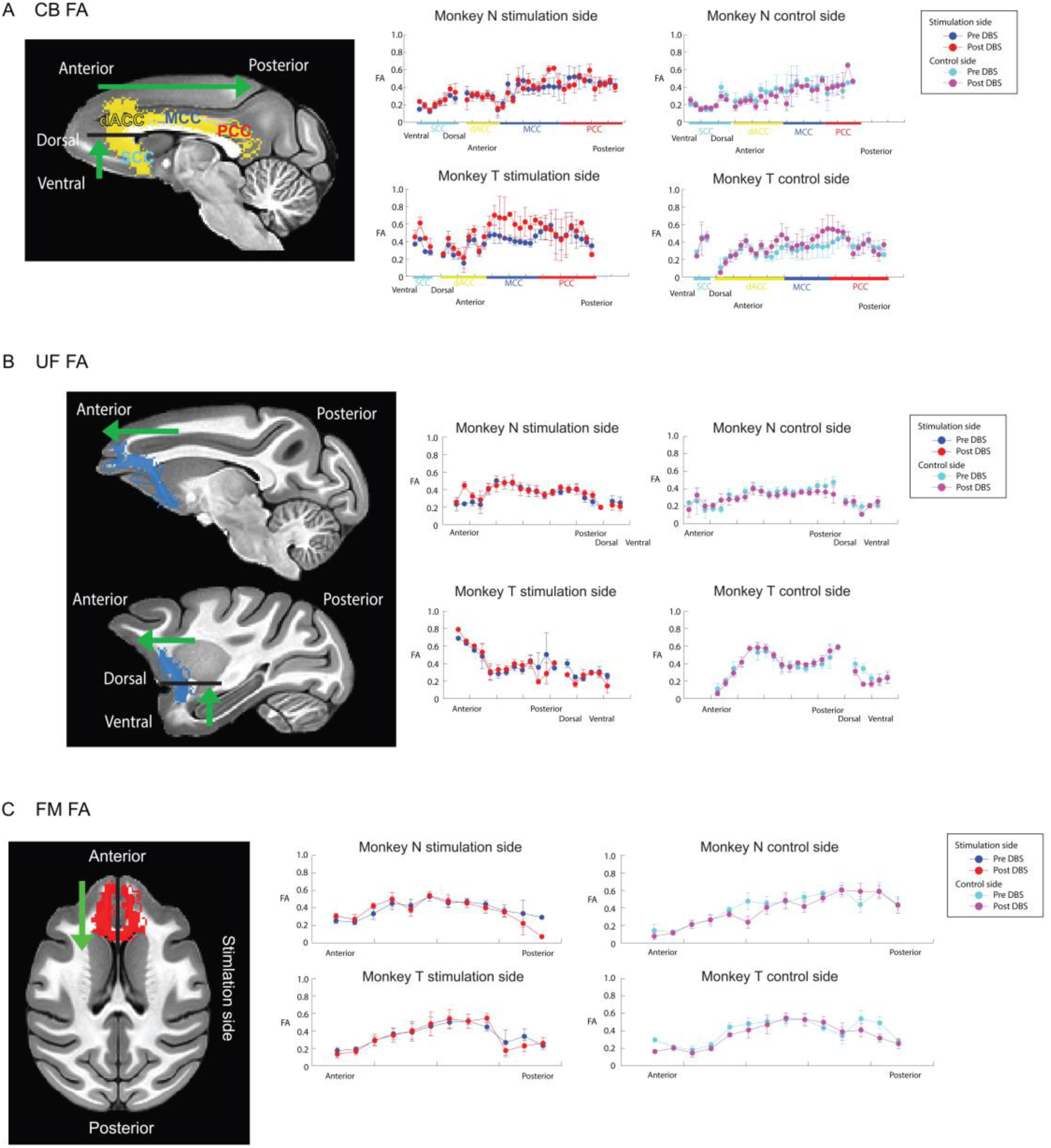
Fine-grained analysis of FA in CB, UF and FM. FA value in CB, UF, and FM extracted from each plane within the tract masks for monkey N (top) and monkey T (bottom). Blue circle/line; pre-DBS averaged FA in the stimulated hemisphere, red circle/line; post-DBS averaged FA in the stimulated hemisphere, cyan circle/line; pre-DBS averaged FA in the control hemisphere, magenta circle/line; post-DBS averaged FA in the control hemisphere. The error bars are SEM. A) The mean FA values per slice in CB. The left image is CB mask (yellow) warped into NMT space from individual pre-DBS T1 space (the sum of monkey N’s CB mask and monkey T’s CB mask). The CB mask was divided into two parts based on its C shape; one part consisted of the SCC, and the other contained the remaining CB regions. For the SCC part, defined as area 25 and area 32 based on the macaque D99 atlas, the mean FA value in each axial slice (the average of FA values extracted from each voxel with the same z coordinates in the individual pre-DBS T1 space) was calculated and plotted. For the remaining part, which includes the dACC, MCC and PCC portion of CB, the mean FA value in each coronal slice (the average of FA values extracted from each voxel which has the same y coordinates) was calculated and plotted. B) The mean FA values per slice in the UF. The left image is UF mask (blue) displayed in NMT space. The UF mask was divided into two parts, the temporal part and the frontal-insula part because of its C shape. For the temporal part, defined as the part of the UF running through the temporal lobe based on macaque D99 atlas, the mean FA value in each axial slice (the average of FA values extracted from each voxel which has the same z coordinates in the individual pre-DBS T1 space) was calculated and plotted in the graphs. For the frontal-insula part, the mean FA in each coronal slice (the average of FA values extracted from each voxel which has the same y coordinates) was calculated and plotted. C) The mean FA values per slice in the FM. The left image is the FM mask (red) displayed in NMT space. The FM mask was divided into left and right hemispheres. For each hemisphere, the mean FA in each coronal slice (the average of FA values extracted from each voxel which has the same y coordinates) was calculated and plotted.

**Supplementary Figure 5:**
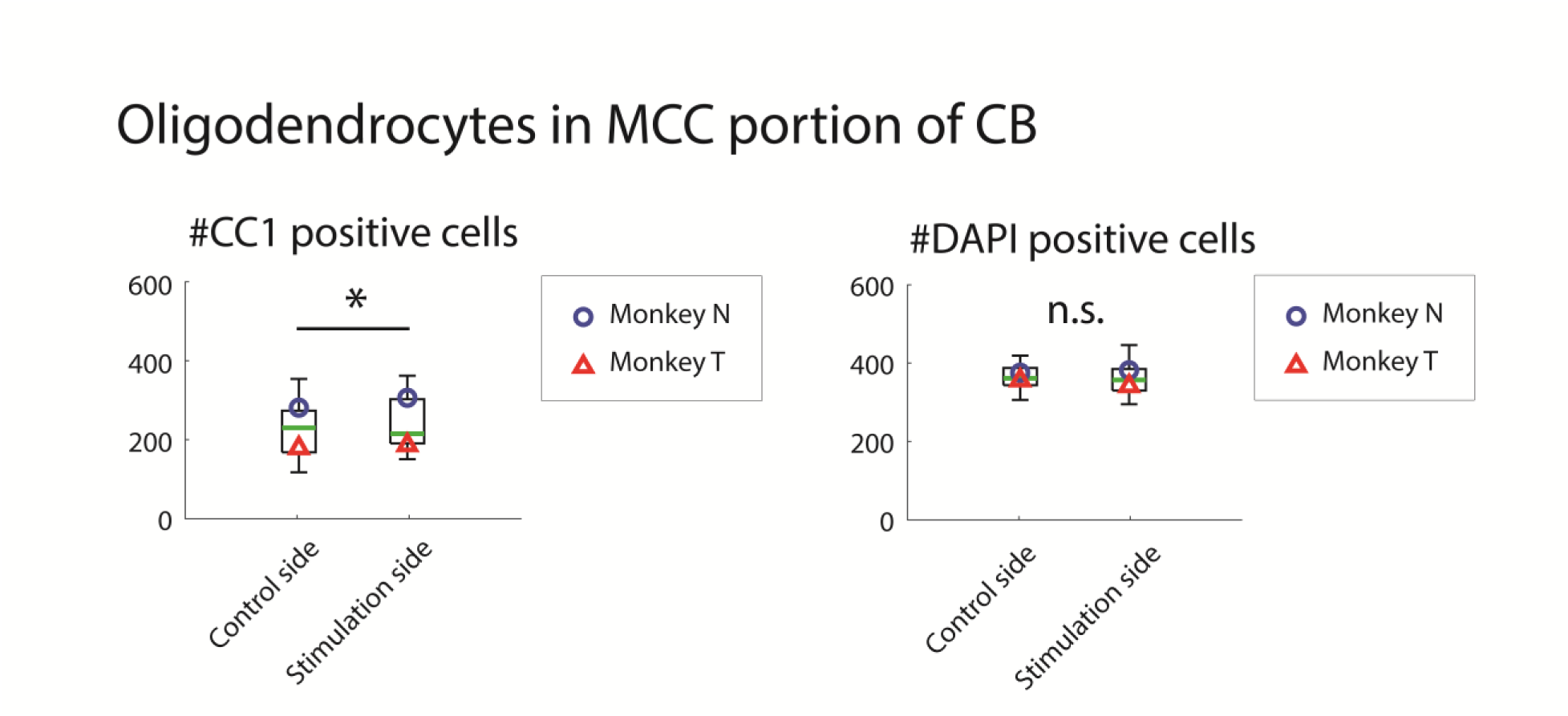
The number of myelinated oligodendrocytes increased in the MCC portion of the CB. Left box plot shows the number of CC1 positive cells (myelinated oligodendrocytes) in the MCC portion of the CB and the right graph is the box plot demonstrating the number of DAPI positive positive cells. Results are shown for both the stimulated hemisphere and the control hemisphere. The green line indicates median values. Symbols represent average values for individual animals (blue circle, monkey N, red triangle, monkey T). One-way repeated measures ANOVA was used (* p < 0.05).

**Supplementary Figure 6:**
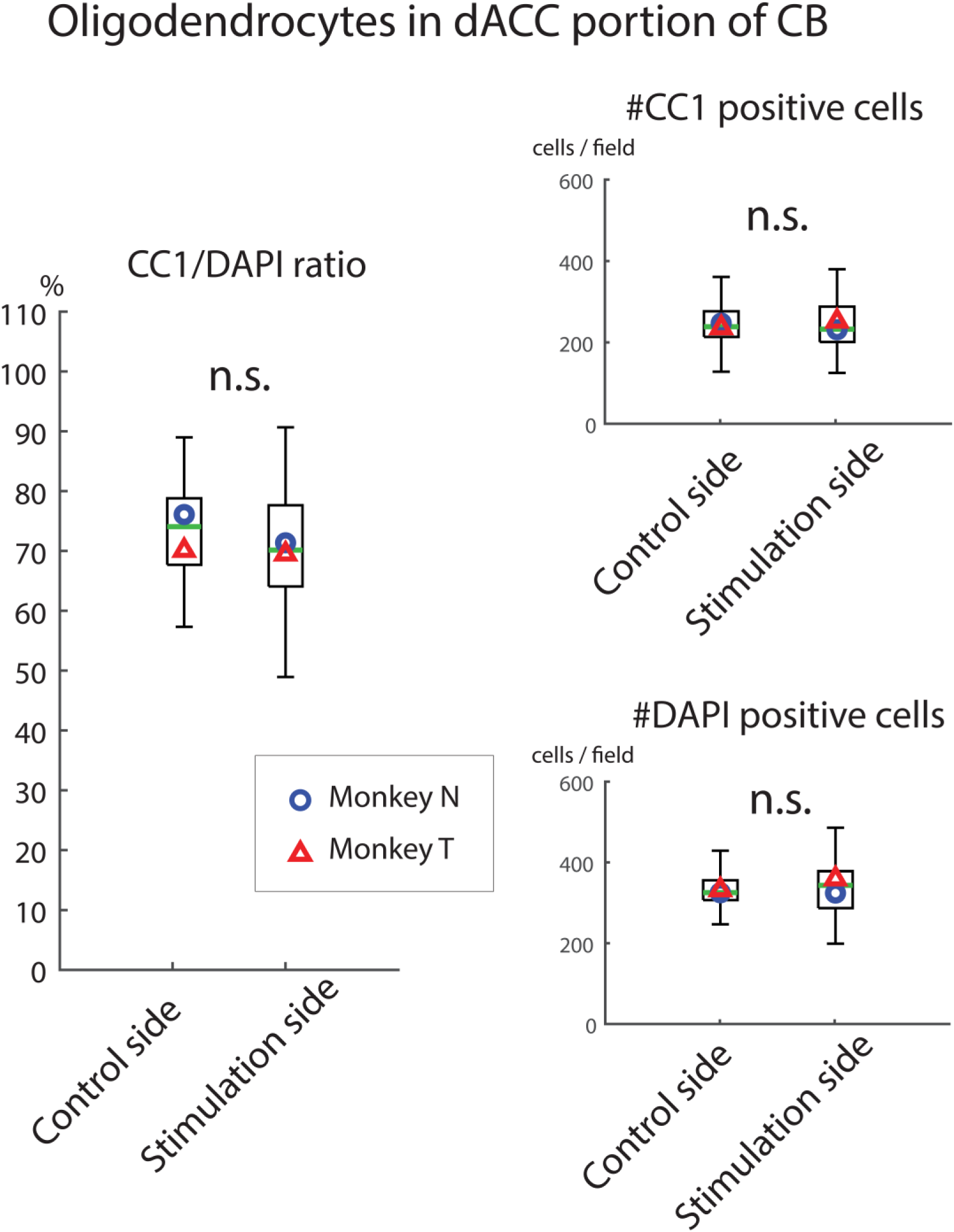
The number of myelinated oligodendrocytes in the dACC portion of the CB. Left box plot shows the ratio of the number of CC1 positive cells to the number of DAPI positive cells in the dACC portion of the CB. The right top graph shows the number of CC1 positive cells and the right bottom graph is a box plot demonstrating the number of DAPI positive cells in the dACC portion of the CB. The green line indicates median values. Symbols represent average values for individual animals (blue circle, monkey N, red triangle, monkey T). One-way repeated measures ANOVA was used (* p < 0.05).

**Supplementary Figure 7:**
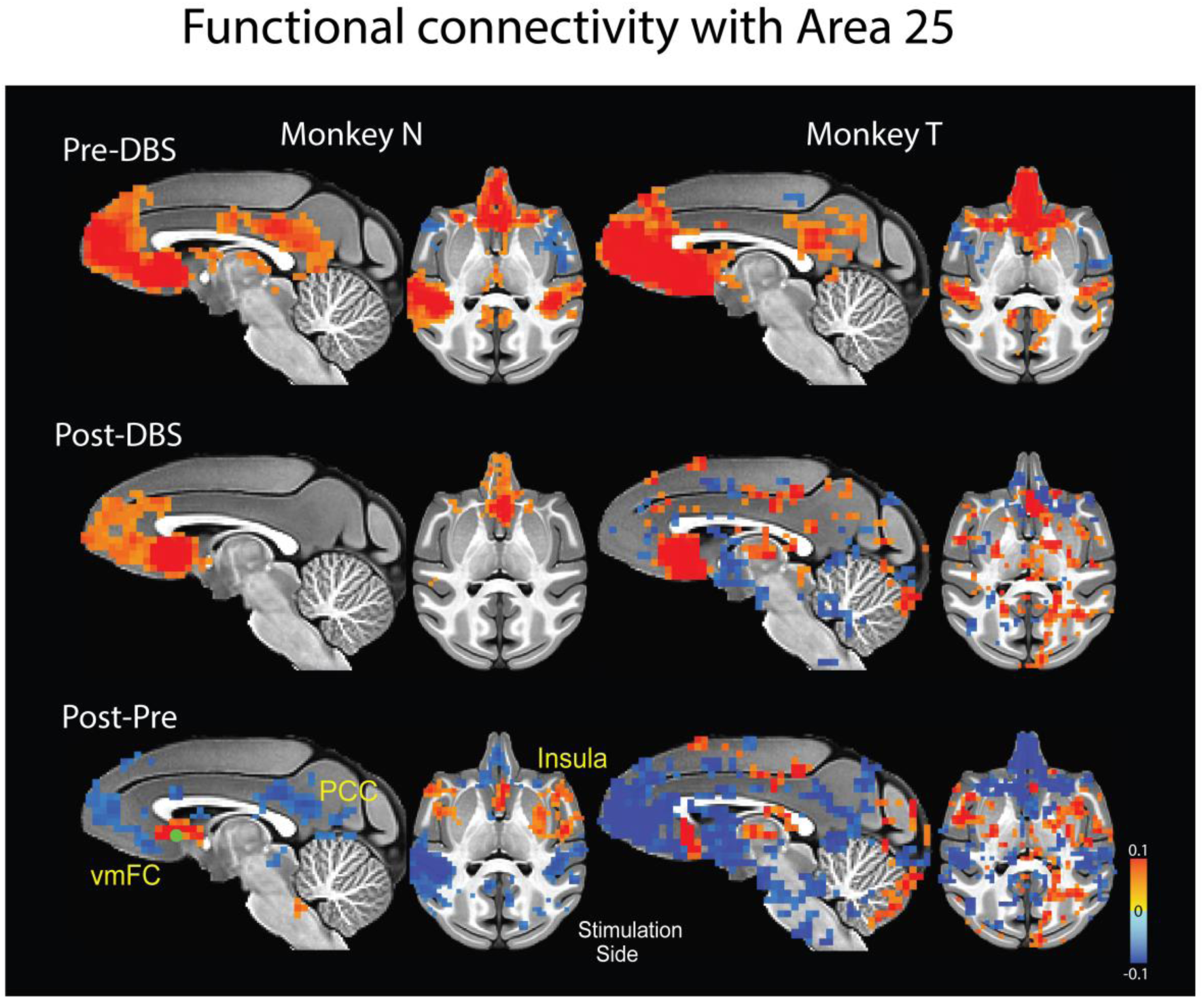
Functional connectivity (FC) with area 25 is altered in individual subjects by SCC-DBS. The functional connectivity (FC) with stimulated area 25 by SCC-DBS of each monkey at pre-DBS (the top row) and post-DBS (the second row) time points. The third row shows the subtraction results of FC with area 25 (post-DBS FC with area 25 – pre-DBS FC with area 25). The left side shows monkey N’s results and the right side showes monkey T’s results. Z-score was calculated in NMT space individually and averaged between the two monkeys after normalizing by the peak z-score in the whole brain. Only FC results that survived thresholding of p < 0.05 and clustering of > 20 voxels are shown. The sagittal plane shows the stimulated hemisphere. In axial images for monkey N, left-right was flipped to show the stimulated hemisphere on the right side of the image. Green circle represents the stimulating contact of the SCC-DBS lead.

**Supplementary Figure 8:**
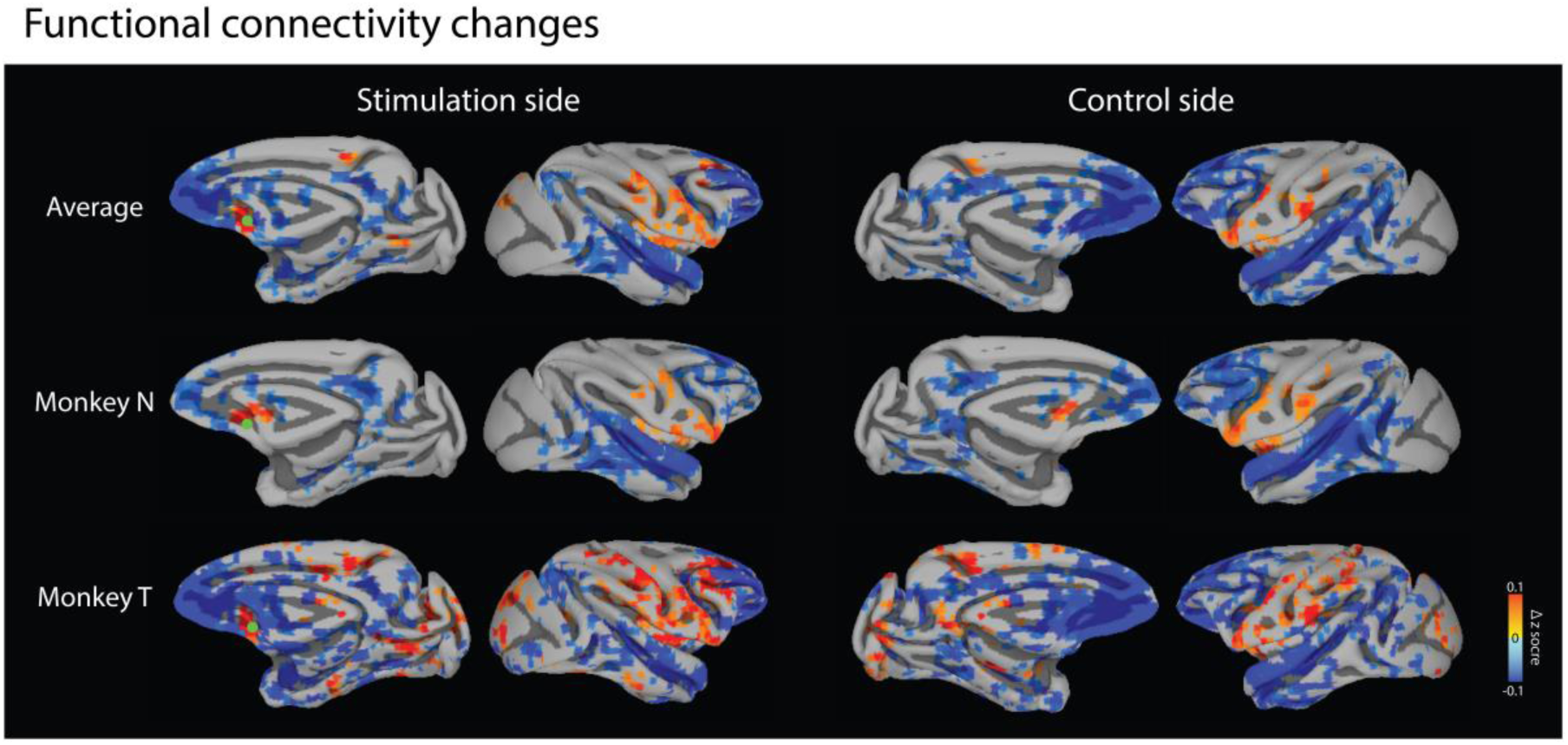
Functional connectivity (FC) with area 25 is altered by SCC-DBS stimulation. The mean functional connectivity (FC) changes with stimulated area 25 by SCC-DBS. Z-score is calculated in NMT space individually and averaged between two monkeys after normalizing by the peak z-score in the whole brain. Only FC results that survived thresholding of p<0.05 and clustering of >20 voxels are shown. All images are the subtraction of pre-stimulation area 25 FC from post-stimulation area 25 FC. The left side shows the stimulated hemisphere and the right side shows the control hemisphere. The first row is the average of 2 monkeys’ FC with stimulated area 25 changes, the second row is the result of monkey N, and the third row is the result of monkey T.

**Supplementary Figure 9:**
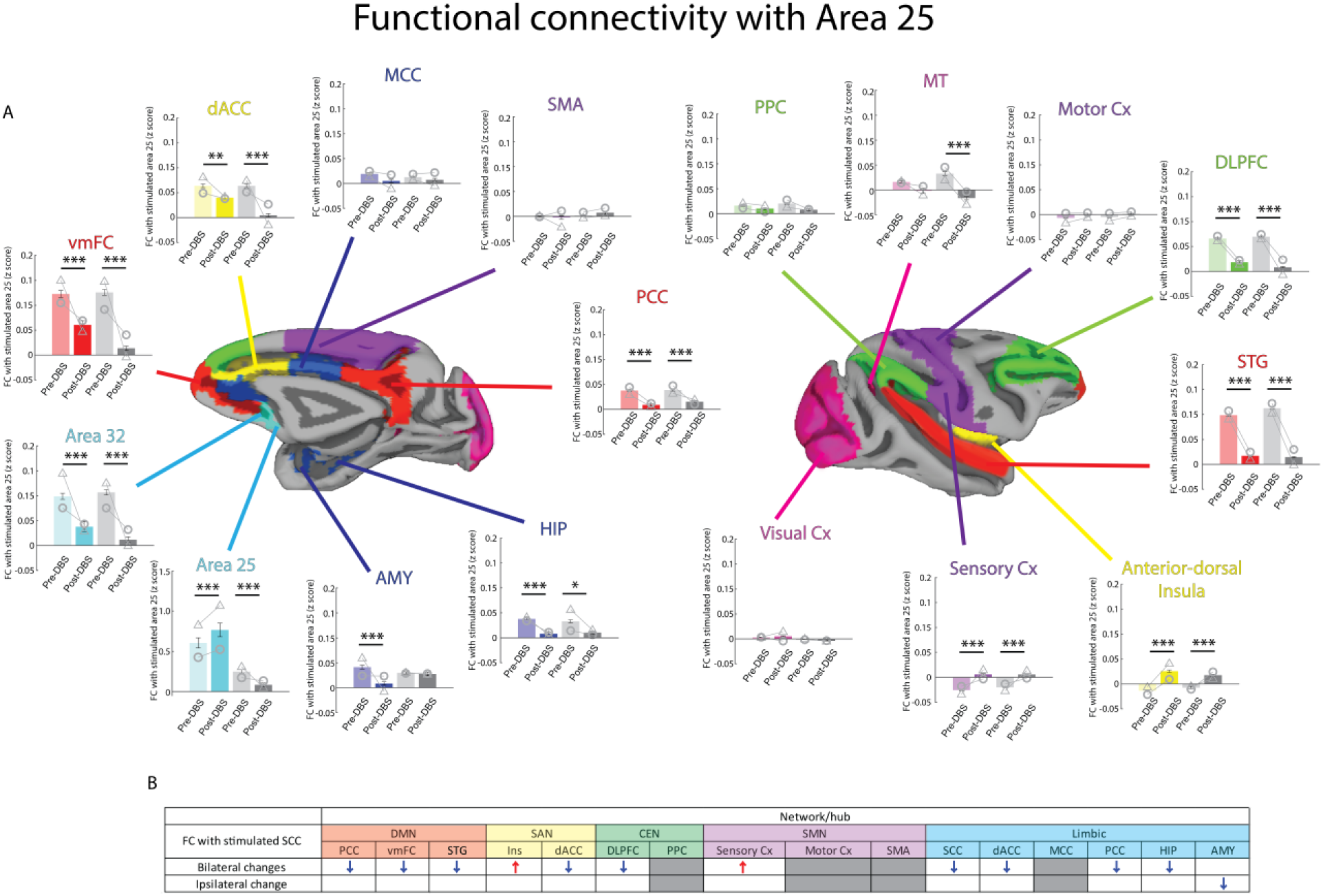
FC with stimulated area 25 changes induced by SCC-DBS in brain network hubs. A) The FC between stimulated area 25 and 17 nodes per hemisphere included in the six brain networks. Colored bars are the mean value of FC normalized z-score between stimulated area 25 and each brain network ROI hub in the stimulated hemisphere (red for the DMN hubs, light blue for SCC (area 25 and area 32), blue for the LIM hubs, yellow for the SAN hubs, light green for the CEN hubs, purple for the SMN hubs and pink for the VIS hubs. SCC, dACC and PCC were counted as hubs for the LIM (blue hubs) as well). These colors are matched with the ROI color in the centered brain image. The gray bars show the mean FC with area 25 in the control hemisphere. Lighter color or lighter gray bars indicate pre-DBS FC z-score with stimulated area 25 and the darker color or darker gray bars represent post-DBS scores. The gray circle represents monkey N and the gray triangle represents monkey T. Error bars show SEM. Three-way repeated measures ANOVA followed by post hoc Turkey-Kramer test were conducted to compare the effects of DBS treatment (* p < 0.05, ** p < 0.01, *** p < 0.001). B) The summary table for the FC between stimulated area 25 and brain network hubs. Significance as calculated by post-hoc analysis (Turkey-Krammer test) on DBS treatment effect (pre-DBS and post-DBS) is indicated by an arrow (p < 0.05). The red arrows indicate significant increases in FC with stimulated area 25 and the blue arrows indicate significant decreases in FC with stimulated area 25.

## References

1. M. Zhdanava, D. Pilon, I. Ghelerter, W. Chow, K. Joshi, P. Lefebvre, J. J. Sheehan, The Prevalence and National Burden of Treatment-Resistant Depression and Major Depressive Disorder in the United States. J Clin Psychiatry 82 (2021).

2. D. J. Lee, C. S. Lozano, R. F. Dallapiazza, A. M. Lozano, Current and future directions of deep brain stimulation for neurological and psychiatric disorders: JNSPG 75th Anniversary Invited Review Article. Journal of Neurosurgery JNS 131, 333–342 (2019).

3. H. S. Mayberg, A. M. Lozano, V. Voon, H. E. McNeely, D. Seminowicz, C. Hamani, J. M. Schwalb, S. H. Kennedy, Deep brain stimulation for treatment-resistant depression. Neuron 45, 651–660 (2005).

4. I. O. Bergfeld, M. Mantione, M. L. C. Hoogendoorn, H. G. Ruhé, P. Notten, J. van Laarhoven, I. Visser, M. Figee, B. P. de Kwaasteniet, F. Horst, A. H. Schene, P. van den Munckhof, G. Beute, R. Schuurman, D. Denys, Deep Brain Stimulation of the Ventral Anterior Limb of the Internal Capsule for Treatment-Resistant Depression: A Randomized Clinical Trial. JAMA Psychiatry 73, 456–464 (2016).

5. A. S. Ríos, S. Oxenford, C. Neudorfer, K. Butenko, N. Li, N. Rajamani, A. Boutet, G. J. B. Elias, J. Germann, A. Loh, W. Deeb, F. Wang, K. Setsompop, B. Salvato, L. B. de Almeida, K. D. Foote, R. Amaral, P. B. Rosenberg, D. F. Tang-Wai, D. A. Wolk, A. D. Burke, S. Salloway, M. N. Sabbagh, M. M. Chakravarty, G. S. Smith, C. G. Lyketsos, M. S. Okun, W. S. Anderson, Z. Mari, F. A. Ponce, A. M. Lozano, A. Horn, Optimal deep brain stimulation sites and networks for stimulation of the fornix in Alzheimer’s disease. Nat Commun 13, 7707 (2022).

6. M. C. Rodriguez-Oroz, J. A. Obeso, A. E. Lang, J.-L. Houeto, P. Pollak, S. Rehncrona, J. Kulisevsky, A. Albanese, J. Volkmann, M. I. Hariz, N. P. Quinn, J. D. Speelman, J. Guridi, I. Zamarbide, A. Gironell, J. Molet, B. Pascual-Sedano, B. Pidoux, A. M. Bonnet, Y. Agid, J. Xie, A.-L. Benabid, A. M. Lozano, J. Saint-Cyr, L. Romito, M. F. Contarino, M. Scerrati, V. Fraix, N. Van Blercom, Bilateral deep brain stimulation in Parkinson’s disease: a multicentre study with 4 years follow-up. Brain 128, 2240–2249 (2005).

7. G. Deuschl, C. Schade-Brittinger, P. Krack, J. Volkmann, H. Schäfer, K. Bötzel, C. Daniels, A. Deutschländer, U. Dillmann, W. Eisner, D. Gruber, W. Hamel, J. Herzog, R. Hilker, S. Klebe, M. Kloss, J. Koy, M. Krause, A. Kupsch, D. Lorenz, S. Lorenzl, H. M. Mehdorn, J. R. Moringlane, W. Oertel, M. O. Pinsker, H. Reichmann, A. Reuss, G.-H. Schneider, A. Schnitzler, U. Steude, V. Sturm, L. Timmermann, V. Tronnier, T. Trottenberg, L. Wojtecki, E. Wolf, W. Poewe, J. Voges, N. S. German Parkinson Study Group, A randomized trial of deep-brain stimulation for Parkinson’s disease. N Engl J Med 355, 896–908 (2006).

8. A. L. Benabid, S. Chabardes, J. Mitrofanis, P. Pollak, Deep brain stimulation of the subthalamic nucleus for the treatment of Parkinson’s disease. Lancet Neurol 8, 67–81 (2009).

9. A. L. Benabid, P. Pollak, C. Gervason, D. Hoffmann, D. M. Gao, M. Hommel, J. E. Perret, J. de Rougemont, Long-term suppression of tremor by chronic stimulation of the ventral intermediate thalamic nucleus. Lancet 337, 403–406 (1991).

10. W. C. Koller, K. E. Lyons, S. B. Wilkinson, A. I. Troster, R. Pahwa, Long-term safety and efficacy of unilateral deep brain stimulation of the thalamus in essential tremor. Mov Disord 16, 464–468 (2001).

11. D. D. Dougherty, A. R. Rezai, L. L. Carpenter, R. H. Howland, M. T. Bhati, J. P. O’Reardon, E. N. Eskandar, G. H. Baltuch, A. D. Machado, D. Kondziolka, C. Cusin, K. C. Evans, L. H. Price, K. Jacobs, M. Pandya, T. Denko, A. R. Tyrka, T. Brelje, T. Deckersbach, C. Kubu, D. A. Malone Jr, A Randomized Sham-Controlled Trial of Deep Brain Stimulation of the Ventral Capsule/Ventral Striatum for Chronic Treatment-Resistant Depression. Biol Psychiatry 78, 240–248 (2014).

12. V. A. Coenen, B. H. Bewernick, S. Kayser, H. Kilian, J. Boström, S. Greschus, R. Hurlemann, M. E. Klein, S. Spanier, B. Sajonz, H. Urbach, T. E. Schlaepfer, Superolateral medial forebrain bundle deep brain stimulation in major depression: a gateway trial. Neuropsychopharmacology 44, 1224–1232 (2019).

13. P. H. Rudebeck, E. L. Rich, H. S. Mayberg, From bed to bench side: Reverse translation to optimize neuromodulation for mood disorders. Proc Natl Acad Sci U S A 116, 26288–26296 (2019).

14. A. M. Lozano, N. Lipsman, H. Bergman, P. Brown, S. Chabardes, J. W. Chang, K. Matthews, C. C. McIntyre, T. E. Schlaepfer, M. Schulder, Y. Temel, J. Volkmann, J. K. Krauss, Deep brain stimulation: current challenges and future directions. Nat Rev Neurol 15, 148–160 (2019).

15. C. Hamani, H. Mayberg, B. Snyder, P. Giacobbe, S. Kennedy, A. M. Lozano, Deep brain stimulation of the subcallosal cingulate gyrus for depression: anatomical location of active contacts in clinical responders and a suggested guideline for targeting: Clinical article. Journal of Neurosurgery JNS 111, 1209–1215 (2009).

16. A. M. Lozano, P. Giacobbe, C. Hamani, S. J. Rizvi, S. H. Kennedy, T. T. Kolivakis, G. Debonnel, A. F. Sadikot, R. W. Lam, A. K. Howard, M. Ilcewicz-Klimek, C. R. Honey, H. S. Mayberg, A multicenter pilot study of subcallosal cingulate area deep brain stimulation for treatment-resistant depression: Clinical article. J Neurosurg 116, 315–322 (2012).

17. P. E. Holtzheimer, M. E. Kelley, R. E. Gross, M. M. Filkowski, S. J. Garlow, A. Barrocas, D. Wint, M. C. Craighead, J. Kozarsky, R. Chismar, J. L. Moreines, K. Mewes, P. R. Posse, D. A. Gutman, H. S. Mayberg, Subcallosal Cingulate Deep Brain Stimulation for Treatment-Resistant Unipolar and Bipolar Depression. Arch Gen Psychiatry 69, 150–158 (2012).

18. H. S. Mayberg, Targeted electrode-based modulation of neural circuits for depression. J Clin Invest 119, 717–725 (2009).

19. H. Johansen-Berg, D. A. Gutman, T. E. J. Behrens, P. M. Matthews, M. F. S. Rushworth, E. Katz, A. M. Lozano, H. S. Mayberg, Anatomical Connectivity of the Subgenual Cingulate Region Targeted with Deep Brain Stimulation for Treatment-Resistant Depression. Cerebral Cortex 18, 1374–1383 (2008).

20. P. Riva-Posse, K. S. Choi, P. E. Holtzheimer, C. C. McIntyre, R. E. Gross, A. Chaturvedi, A. L. Crowell, S. J. Garlow, J. K. Rajendra, H. S. Mayberg, Defining critical white matter pathways mediating successful subcallosal cingulate deep brain stimulation for treatment-resistant depression. Biol Psychiatry 76, 963– 969 (2014).

21. K. S. Choi, P. Riva-Posse, R. E. Gross, H. S. Mayberg, Mapping the “Depression Switch” During Intraoperative Testing of Subcallosal Cingulate Deep Brain Stimulation. JAMA Neurol 72, 1252–1260 (2015).

22. P. Riva-Posse, K. S. Choi, P. E. Holtzheimer, A. L. Crowell, S. J. Garlow, J. K. Rajendra, C. C. McIntyre, R. E. Gross, H. S. Mayberg, A connectomic approach for subcallosal cingulate deep brain stimulation surgery: prospective targeting in treatment-resistant depression. Mol Psychiatry 23, 843–849 (2018).

23. G. J. B. Elias, J. Germann, A. Boutet, M. E. Beyn, P. Giacobbe, H. N. Song, K. S. Choi, H. S. Mayberg, S. H. Kennedy, A. M. Lozano, Local neuroanatomical and tract-based proxies of optimal subcallosal cingulate deep brain stimulation. Brain Stimul 16, 1259–1272 (2023).

24. S. H. Kennedy, P. Giacobbe, S. J. Rizvi, F. M. Placenza, Y. Nishikawa, H. S. Mayberg, A. M. Lozano, Deep brain stimulation for treatment-resistant depression: follow-up after 3 to 6 years. Am J Psychiatry 168, 502–510 (2011).

25. A. L. Crowell, P. Riva-Posse, P. E. Holtzheimer, S. J. Garlow, M. E. Kelley, R. E. Gross, L. Denison, S. Quinn, H. S. Mayberg, Long-Term Outcomes of Subcallosal Cingulate Deep Brain Stimulation for Treatment-Resistant Depression. Am J Psychiatry 176, 949–956 (2019).

26. K. A. Johnson, M. S. Okun, K. W. Scangos, H. S. Mayberg, C. de Hemptinne, Deep brain stimulation for refractory major depressive disorder: a comprehensive review. Mol Psychiatry, doi: 10.1038/s41380-023-02394-4 (2024).

27. K. J. Ressler, H. S. Mayberg, Targeting abnormal neural circuits in mood and anxiety disorders: from the laboratory to the clinic. Nat Neurosci 10, 1116–1124 (2007).

28. A. M. Lozano, H. S. Mayberg, P. Giacobbe, C. Hamani, R. C. Craddock, S. H. Kennedy, Subcallosal cingulate gyrus deep brain stimulation for treatment-resistant depression. Biol Psychiatry 64, 461–467 (2008).

29. J. M. Broadway, P. E. Holtzheimer, M. R. Hilimire, N. A. Parks, J. E. DeVylder, H. S. Mayberg, P. M. Corballis, Frontal Theta Cordance Predicts 6-Month Antidepressant Response to Subcallosal Cingulate Deep Brain Stimulation for Treatment-Resistant Depression: A Pilot Study. Neuropsychopharmacology 37, 1764–1772 (2012).

30. J. Cha, K. S. Choi, J. K. Rajendra, C. L. McGrath, P. Riva-Posse, P. E. Holtzheimer, M. Figee, B. H. Kopell, H. S. Mayberg, Whole brain network effects of subcallosal cingulate deep brain stimulation for treatment-resistant depression. Mol Psychiatry 29, 112–120 (2024).

31. M. S. E. Sendi, A. C. Waters, V. Tiruvadi, P. Riva-Posse, A. Crowell, F. Isbaine, J. T. Gale, K. S. Choi, R. E. Gross, H. S. Mayberg, B. Mahmoudi, Intraoperative neural signals predict rapid antidepressant effects of deep brain stimulation. Transl Psychiatry 11, 551 (2021).

32. S. Alagapan, K. S. Choi, S. Heisig, P. Riva-Posse, A. Crowell, V. Tiruvadi, M. Obatusin, A. Veerakumar, A. C. Waters, R. E. Gross, S. Quinn, L. Denison, M. O’Shaughnessy, M. Connor, G. Canal, J. Cha, R. Hershenberg, T. Nauvel, F. Isbaine, M. F. Afzal, M. Figee, B. H. Kopell, R. Butera, H. S. Mayberg, C. J. Rozell, Cingulate dynamics track depression recovery with deep brain stimulation. Nature 622, 130–138 (2023).

33. G. J. B. Elias, J. Germann, A. Boutet, A. Pancholi, M. E. Beyn, K. Bhatia, C. Neudorfer, A. Loh, S. J. Rizvi, V. Bhat, P. Giacobbe, D. B. Woodside, S. H. Kennedy, A. M. Lozano, Structuro-functional surrogates of response to subcallosal cingulate deep brain stimulation for depression. Brain 145, 362–377 (2022).

34. E. E. Smith, K. S. Choi, A. Veerakumar, M. Obatusin, B. Howell, A. H. Smith, V. Tiruvadi, A. L. Crowell, P. Riva-Posse, S. Alagapan, C. J. Rozell, H. S. Mayberg, A. C. Waters, Time-frequency signatures evoked by single-pulse deep brain stimulation to the subcallosal cingulate. Front Hum Neurosci 16, 939258 (2022).

35. R. J. Zatorre, R. D. Fields, H. Johansen-Berg, Plasticity in gray and white: neuroimaging changes in brain structure during learning. Nat Neurosci 15, 528–536 (2012).

36. H. Johansen-Berg, Behavioural relevance of variation in white matter microstructure. Curr Opin Neurol 23, 351–358 (2010).

37. S. Chanraud, N. Zahr, E. V Sullivan, A. Pfefferbaum, MR Diffusion Tensor Imaging: A Window into White Matter Integrity of the Working Brain. Neuropsychol Rev 20, 209–225 (2010).

38. S.-K. Song, S.-W. Sun, M. J. Ramsbottom, C. Chang, J. Russell, A. H. Cross, Dysmyelination Revealed through MRI as Increased Radial (but Unchanged Axial) Diffusion of Water. Neuroimage 17, 1429–1436 (2002).

39. A. Seehaus, A. Roebroeck, M. Bastiani, L. Fonseca, H. Bratzke, N. Lori, A. Vilanova, R. Goebel, R. Galuske, Histological validation of high-resolution DTI in human post mortem tissue. Front Neuroanat 9, 98 (2015).

40. S. R. Heilbronner, S. N. Haber, Frontal Cortical and Subcortical Projections Provide a Basis for Segmenting the Cingulum Bundle: Implications for Neuroimaging and Psychiatric Disorders. The Journal of Neuroscience 34, 10041 (2014).

41. E. J. Bubb, C. Metzler-Baddeley, J. P. Aggleton, The cingulum bundle: Anatomy, function, and dysfunction. Neurosci Biobehav Rev 92, 104–127 (2018).

42. Y. Wu, D. Sun, Y. Wang, Y. Wang, S. Ou, Segmentation of the Cingulum Bundle in the Human Brain: A New Perspective Based on DSI Tractography and Fiber Dissection Study. Front Neuroanat 10, 84 (2016).

43. P. Friedrich, C. Fraenz, C. Schlüter, S. Ocklenburg, B. Mädler, O. Güntürkün, E. Genç, The Relationship Between Axon Density, Myelination, and Fractional Anisotropy in the Human Corpus Callosum. Cerebral Cortex 30, 2042–2056 (2020).

44. C. Beaulieu, The basis of anisotropic water diffusion in the nervous system - a technical review. NMR Biomed 15, 435–455 (2002).

45. S. Boretius, A. Escher, T. Dallenga, C. Wrzos, R. Tammer, W. Brück, S. Nessler, J. Frahm, C. Stadelmann, Assessment of lesion pathology in a new animal model of MS by multiparametric MRI and DTI. Neuroimage 59, 2678–2688 (2012).

46. R. D. Fields, A new mechanism of nervous system plasticity: activity-dependent myelination. Nat Rev Neurosci 16, 756–767 (2015).

47. G. Bonetto, D. Belin, R. T. Káradóttir, Myelin: A gatekeeper of activity-dependent circuit plasticity? Science (1979) 374, eaba6905 (2024).

48. F. Kimura, C. Itami, Myelination and isochronicity in neural networks. Front Neuroanat 3, 12 (2009).

49. S. Pajevic, P. J. Basser, R. D. Fields, Role of myelin plasticity in oscillations and synchrony of neuronal activity. Neuroscience 276, 135–147 (2013).

50. S. Pajevic, D. Plenz, P. J. Basser, R. D. Fields, Oligodendrocyte-mediated myelin plasticity and its role in neural synchronization. Elife 12 (2023).

51. B. Emery, Regulation of oligodendrocyte differentiation and myelination. Science (1979) 330, 779–782 (2010).

52. P. Fotiadis, M. Cieslak, X. He, L. Caciagli, M. Ouellet, T. D. Satterthwaite, R. T. Shinohara, D. S. Bassett, Myelination and excitation-inhibition balance synergistically shape structure-function coupling across the human cortex. Nat Commun 14, 6115 (2023).

53. C. J. Honey, O. Sporns, L. Cammoun, X. Gigandet, J. P. Thiran, R. Meuli, P. Hagmann, Predicting human resting-state functional connectivity from structural connectivity. Proc Natl Acad Sci U S A 106, 2035–2040 (2009).

54. A. C. Waters, A. Veerakumar, K. S. Choi, B. Howell, V. Tiruvadi, K. R. Bijanki, A. Crowell, P. Riva-Posse, H. S. Mayberg, Test-retest reliability of a stimulation-locked evoked response to deep brain stimulation in subcallosal cingulate for treatment resistant depression. Hum Brain Mapp 39, 4844–4856 (2018).

55. A. Seas, M. S. Noor, K. S. Choi, A. Veerakumar, M. Obatusin, J. Dahill-Fuchel, V. Tiruvadi, E. Xu, P. Riva-Posse, C. J. Rozell, H. S. Mayberg, C. C. McIntyre, A. C. Waters, B. Howell, Subcallosal cingulate deep brain stimulation evokes two distinct cortical responses via differential white matter activation. Proceedings of the National Academy of Sciences 121, e2314918121 (2024).

56. P. N. Alves, C. Foulon, V. Karolis, D. Bzdok, D. S. Margulies, E. Volle, M. Thiebaut de Schotten, An improved neuroanatomical model of the default-mode network reconciles previous neuroimaging and neuropathological findings. Commun Biol 2, 370 (2019).

57. M. van den Heuvel, R. Mandl, J. Luigjes, H. Hulshoff Pol, Microstructural Organization of the Cingulum Tract and the Level of Default Mode Functional Connectivity. The Journal of Neuroscience 28, 10844 (2008).

58. P. Qin, G. Northoff, How is our self related to midline regions and the default-mode network? Neuroimage 57, 1221–1233 (2011).

59. J. P. Hamilton, M. Farmer, P. Fogelman, I. H. Gotlib, Depressive Rumination, the Default-Mode Network, and the Dark Matter of Clinical Neuroscience. Biol Psychiatry 78, 224–230 (2015).

60. A. T. Drysdale, L. Grosenick, J. Downar, K. Dunlop, F. Mansouri, Y. Meng, R. N. Fetcho, B. Zebley, D. J. Oathes, A. Etkin, A. F. Schatzberg, K. Sudheimer, J. Keller, H. S. Mayberg, F. M. Gunning, G. S. Alexopoulos, M. D. Fox, A. Pascual-Leone, H. U. Voss, B. J. Casey, M. J. Dubin, C. Liston, Resting-state connectivity biomarkers define neurophysiological subtypes of depression. Nat Med 23, 28–38 (2017).

61. A. Scalabrini, B. Vai, S. Poletti, S. Damiani, C. Mucci, C. Colombo, R. Zanardi, F. Benedetti, G. Northoff, All roads lead to the default-mode network—global source of DMN abnormalities in major depressive disorder. Neuropsychopharmacology 45, 2058–2069 (2020).

62. B. J. Borserio, C. F. Sharpley, V. Bitsika, K. Sarmukadam, P. J. Fourie, L. L. Agnew, Default mode network activity in depression subtypes. 32, 597–613 (2021).

63. N. Javaheripour, L. Colic, N. Opel, M. Li, S. Maleki Balajoo, T. Chand, J. Van der Meer, M. Krylova, I. Izyurov, T. Meller, J. Goltermann, N. R. Winter, S. Meinert, D. Grotegerd, A. Jansen, N. Alexander, P. Usemann, F. Thomas-Odenthal, U. Evermann, A. Wroblewski, K. Brosch, F. Stein, T. Hahn, B. Straube, A. Krug, I. Nenadić, T. Kircher, I. Croy, U. Dannlowski, G. Wagner, M. Walter, Altered brain dynamic in major depressive disorder: state and trait features. Transl Psychiatry 13, 261 (2023).

64. D. Fennema, G. J. Barker, O. O’Daly, S. Duan, E. Carr, K. Goldsmith, A. H. Young, J. Moll, R. Zahn, The Role of Subgenual Resting-State Connectivity Networks in Predicting Prognosis in Major Depressive Disorder. Biological Psychiatry Global Open Science 4, 100308 (2024).

65. C.-C. Huang, Q. Luo, L. Palaniyappan, A. C. Yang, C.-C. Hung, K.-H. Chou, C.-Y. Zac Lo, M.-N. Liu, S.-J. Tsai, D. M. Barch, J. Feng, C.-P. Lin, T. W. Robbins, Transdiagnostic and Illness-Specific Functional Dysconnectivity Across Schizophrenia, Bipolar Disorder, and Major Depressive Disorder. Biol Psychiatry Cogn Neurosci Neuroimaging 5, 542–553 (2020).

66. A. Lazari, P. Salvan, L. Verhagen, M. Cottaar, D. Papp, O. J. van der Werf, B. Gavine, J. Kolasinski, M. Webster, C. J. Stagg, M. F. S. Rushworth, H. Johansen-Berg, A macroscopic link between interhemispheric tract myelination and cortico-cortical interactions during action reprogramming. Nat Commun 13, 4253 (2022).

67. E. M. Gibson, D. Purger, C. W. Mount, A. K. Goldstein, G. L. Lin, L. S. Wood, I. Inema, S. E. Miller, G. Bieri, J. B. Zuchero, B. A. Barres, P. J. Woo, H. Vogel, M. Monje, Neuronal Activity Promotes Oligodendrogenesis and Adaptive Myelination in the Mammalian Brain. Science (1979) 344, 1252304 (2014).

68. S. Mitew, I. Gobius, L. R. Fenlon, S. J. McDougall, D. Hawkes, Y. L. Xing, H. Bujalka, A. L. Gundlach, L. J. Richards, T. J. Kilpatrick, T. D. Merson, B. Emery, Pharmacogenetic stimulation of neuronal activity increases myelination in an axon-specific manner. Nat Commun 9, 306 (2018).

69. D. Ongür, W. C. Drevets, J. L. Price, Glial reduction in the subgenual prefrontal cortex in mood disorders. Proc Natl Acad Sci U S A 95, 13290–13295 (1998).

70. G. Rajkowska, J. J. Miguel-Hidalgo, J. Wei, G. Dilley, S. D. Pittman, H. Y. Meltzer, J. C. Overholser, B. L. Roth, C. A. Stockmeier, Morphometric evidence for neuronal and glial prefrontal cell pathology in major depression. Biol Psychiatry 45, 1085–1098 (1999).

71. T. Bracht, D. Linden, P. Keedwell, A review of white matter microstructure alterations of pathways of the reward circuit in depression. J Affect Disord 187, 45–53 (2015).

72. D. M. Barch, X. Hua, S. Kandala, M. P. Harms, A. Sanders, R. Brady, R. Tillman, J. L. Luby, White matter alterations associated with lifetime and current depression in adolescents: Evidence for cingulum disruptions. Depress Anxiety 39, 881–890 (2022).

73. L. S. van Velzen, S. Kelly, D. Isaev, A. Aleman, L. I. Aftanas, J. Bauer, B. T. Baune, I. V Brak, A. Carballedo, C. G. Connolly, B. Couvy-Duchesne, K. R. Cullen, K. V Danilenko, U. Dannlowski, V. Enneking, E. Filimonova, K. Förster, T. Frodl, I. H. Gotlib, N. A. Groenewold, D. Grotegerd, M. A. Harris, S. N. Hatton, E. L. Hawkins, I. B. Hickie, T. C. Ho, A. Jansen, T. Kircher, B. Klimes-Dougan, P. Kochunov, A. Krug, J. Lagopoulos, R. Lee, T. A. Lett, M. Li, F. P. MacMaster, N. G. Martin, A. M. McIntosh, Q. McLellan, S. Meinert, I. Nenadić, E. Osipov, B. W. J. H. Penninx, M. J. Portella, J. Repple, A. Roos, M. D. Sacchet, P. G. Sämann, K. Schnell, X. Shen, K. Sim, D. J. Stein, M.-J. van Tol, A. S. Tomyshev, L. Tozzi, I. M. Veer, R. Vermeiren, Y. Vives-Gilabert, H. Walter, M. Walter, N. J. A. van der Wee, S. J. A. van der Werff, M. W. Schreiner, H. C. Whalley, M. J. Wright, T. T. Yang, A. Zhu, D. J. Veltman, P. M. Thompson, N. Jahanshad, L. Schmaal, White matter disturbances in major depressive disorder: a coordinated analysis across 20 international cohorts in the ENIGMA MDD working group. Mol Psychiatry 25, 1511–1525 (2020).

74. V. J. Sydnor, A. E. Lyall, S. Cetin-Karayumak, J. C. Cheung, J. M. Felicione, O. Akeju, M. E. Shenton, T. Deckersbach, D. F. Ionescu, O. Pasternak, C. Cusin, M. Kubicki, Studying pre-treatment and ketamine-induced changes in white matter microstructure in the context of ketamine’s antidepressant effects. Transl Psychiatry 10, 432 (2020).

75. M. M. Vasavada, A. M. Leaver, R. T. Espinoza, S. H. Joshi, S. N. Njau, R. P. Woods, K. L. Narr, Structural connectivity and response to ketamine therapy in major depression: A preliminary study. J Affect Disord 190, 836–841 (2016).

76. G. Deco, V. K. Jirsa, A. R. McIntosh, Emerging concepts for the dynamical organization of resting-state activity in the brain. Nat Rev Neurosci 12, 43–56 (2011).

77. M. L. Kringelbach, A. L. Green, T. Z. Aziz, Balancing the brain: resting state networks and deep brain stimulation. Front Integr Neurosci 5, 8 (2011).

78. D. Vidaurre, R. Abeysuriya, R. Becker, A. J. Quinn, F. Alfaro-Almagro, S. M. Smith, M. W. Woolrich, Discovering dynamic brain networks from big data in rest and task. Neuroimage 180, 646–656 (2018).

79. D. Lyu, S. Naik, D. K. Menon, E. A. Stamatakis, Intrinsic brain dynamics in the Default Mode Network predict involuntary fluctuations of visual awareness. Nat Commun 13, 6923 (2022).

80. O. Smart, K. S. Choi, P. Riva-Posse, V. Tiruvadi, J. Rajendra, A. C. Waters, A. L. Crowell, J. Edwards, R. E. Gross, H. S. Mayberg, Initial Unilateral Exposure to Deep Brain Stimulation in Treatment-Resistant Depression Patients Alters Spectral Power in the Subcallosal Cingulate. Front Comput Neurosci 12, 43 (2018).

81. R. G. Almeida, D. A. Lyons, On Myelinated Axon Plasticity and Neuronal Circuit Formation and Function. The Journal of Neuroscience 37, 10023 (2017).

82. E. G. Hughes, J. L. Orthmann-Murphy, A. J. Langseth, D. E. Bergles, Myelin remodeling through experience-dependent oligodendrogenesis in the adult somatosensory cortex. Nat Neurosci 21, 696–706 (2018).

83. M. P. van den Heuvel, O. Sporns, Network hubs in the human brain. Trends Cogn Sci 17, 683–696 (2013).

84. I. R. Kleckner, J. Zhang, A. Touroutoglou, L. Chanes, C. Xia, W. K. Simmons, K. S. Quigley, B. C. Dickerson, L. F. Barrett, Evidence for a Large-Scale Brain System Supporting Allostasis and Interoception in Humans. Nat Hum Behav 1 (2017).

85. A. Fujimoto, C. Elorette, J. M. Fredericks, S. H. Fujimoto, L. Fleysher, P. H. Rudebeck, B. E. Russ, Resting-State fMRI-Based Screening of Deschloroclozapine in Rhesus Macaques Predicts Dosage-Dependent Behavioral Effects. J Neurosci 42, 5705–5716 (2022).

86. R. M. Hutchison, T. Womelsdorf, J. S. Gati, S. Everling, R. S. Menon, Resting-state networks show dynamic functional connectivity in awake humans and anesthetized macaques. Hum Brain Mapp 34, 2154–2177 (2012).

87. T.-L. Wu, A. Mishra, F. Wang, P.-F. Yang, J. C. Gore, L. M. Chen, Effects of isoflurane anesthesia on resting-state fMRI signals and functional connectivity within primary somatosensory cortex of monkeys. Brain Behav 6, e00591 (2016).

88. C. Giacometti, A. Dureux, D. Autran-Clavagnier, C. R. E. Wilson, J. Sallet, M. Dirheimer, E. Procyk, F. Hadj-Bouziane, C. Amiez, Frontal cortical functional connectivity is impacted by anaesthesia in macaques. Cereb Cortex 32, 4050–4067 (2022).

89. C. Elorette, A. Fujimoto, F. M. Stoll, S. H. Fujimoto, N. Bienkowska, L. London, L. Fleysher, B. E. Russ, P. H. Rudebeck, The neural basis of resting-state fMRI functional connectivity in fronto-limbic circuits revealed by chemogenetic manipulation. Nat Commun 15, 4669 (2024).

90. F. P. Leite, D. Tsao, W. Vanduffel, D. Fize, Y. Sasaki, L. L. Wald, A. M. Dale, K. K. Kwong, G. A. Orban, B. R. Rosen, R. B. H. Tootell, J. B. Mandeville, Repeated fMRI using iron oxide contrast agent in awake, behaving macaques at 3 Tesla. Neuroimage 16, 283–294 (2002).

91. B. E. Russ, C. I. Petkov, S. C. Kwok, Q. Zhu, P. Belin, W. Vanduffel, S. Ben Hamed, Common functional localizers to enhance NHP & cross-species neuroscience imaging research. Neuroimage 237, 118203 (2021).

92. R. W. Cox, AFNI: software for analysis and visualization of functional magnetic resonance neuroimages. Comput. Biomed. Res. 29, 162–173 (1996).

93. M. Jenkinson, C. F. Beckmann, T. E. J. Behrens, M. W. Woolrich, S. M. Smith, FSL. Neuroimage 62, 782–790 (2012).

94. M. W. Woolrich, S. Jbabdi, B. Patenaude, M. Chappell, S. Makni, T. Behrens, C. Beckmann, M. Jenkinson, S. M. Smith, Bayesian analysis of neuroimaging data in FSL. Neuroimage 45, S173–S186 (2009).

95. S. M. Smith, M. Jenkinson, M. W. Woolrich, C. F. Beckmann, T. E. J. Behrens, H. Johansen-Berg, P. R. Bannister, M. De Luca, I. Drobnjak, D. E. Flitney, R. K. Niazy, J. Saunders, J. Vickers, Y. Zhang, N. De Stefano, J. M. Brady, P. M. Matthews, Advances in functional and structural MR image analysis and implementation as FSL. Neuroimage 23, S208–S219 (2004).

96. B. B. Avants, N. J. Tustison, G. Song, P. A. Cook, A. Klein, J. C. Gee, A reproducible evaluation of ANTs similarity metric performance in brain image registration. Neuroimage 54, 2033–2044 (2011).

97. A. Klein, J. Andersson, B. A. Ardekani, J. Ashburner, B. Avants, M.-C. Chiang, G. E. Christensen, D. L. Collins, J. Gee, P. Hellier, J. H. Song, M. Jenkinson, C. Lepage, D. Rueckert, P. Thompson, T. Vercauteren, R. P. Woods, J. J. Mann, R. V Parsey, Evaluation of 14 nonlinear deformation algorithms applied to human brain MRI registration. Neuroimage 46, 786–802 (2009).

98. B. B. Avants, C. L. Epstein, M. Grossman, J. C. Gee, Symmetric diffeomorphic image registration with cross-correlation: Evaluating automated labeling of elderly and neurodegenerative brain. Med Image Anal 12, 26–41 (2008).

99. P. A. Yushkevich, J. Piven, H. C. Hazlett, R. G. Smith, S. Ho, J. C. Gee, G. Gerig, User-guided 3D active contour segmentation of anatomical structures: Significantly improved efficiency and reliability. Neuroimage 31, 1116–1128 (2006).

100. J. Veraart, E. Fieremans, D. S. Novikov, Diffusion MRI noise mapping using random matrix theory. Magn Reson Med 76, 1582–1593 (2016).

101. J. Veraart, D. S. Novikov, D. Christiaens, B. Ades-aron, J. Sijbers, E. Fieremans, Denoising of diffusion MRI using random matrix theory. Neuroimage 142, 394–406 (2016).

102. L. Cordero-Grande, D. Christiaens, J. Hutter, A. N. Price, J. V Hajnal, Complex diffusion-weighted image estimation via matrix recovery under general noise models. Neuroimage 200, 391–404 (2019).

103. J.-D. Tournier, R. Smith, D. Raffelt, R. Tabbara, T. Dhollander, M. Pietsch, D. Christiaens, B. Jeurissen, C.-H. Yeh, A. Connelly, MRtrix3: A fast, flexible and open software framework for medical image processing and visualisation. Neuroimage 202, 116137 (2019).

104. N. J. Tustison, B. B. Avants, P. A. Cook, Y. Zheng, A. Egan, P. A. Yushkevich, J. C. Gee, N4ITK: Improved N3 Bias Correction. IEEE Trans Med Imaging 29, 1310–1320 (2010).

105. T. E. J. Behrens, M. W. Woolrich, M. Jenkinson, H. Johansen-Berg, R. G. Nunes, S. Clare, P. M. Matthews, J. M. Brady, S. M. Smith, Characterization and propagation of uncertainty in diffusion-weighted MR imaging. Magn Reson Med 50, 1077–1088 (2003).

106. T. E. J. Behrens, H. J. Berg, S. Jbabdi, M. F. S. Rushworth, M. W. Woolrich, Probabilistic diffusion tractography with multiple fibre orientations: What can we gain? Neuroimage 34, 144–155 (2007).

107. D. Folloni, J. Sallet, A. A. Khrapitchev, N. Sibson, L. Verhagen, R. B. Mars, Dichotomous organization of amygdala/temporal-prefrontal bundles in both humans and monkeys. Elife 8, e47175 (2019).

108. S. Warrington, K. L. Bryant, A. A. Khrapitchev, J. Sallet, M. Charquero-Ballester, G. Douaud, S. Jbabdi, R. B. Mars, S. N. Sotiropoulos, XTRACT - Standardised protocols for automated tractography in the human and macaque brain. Neuroimage 217, 116923 (2020).

109. J. D. Schmahmann, D. N. Pandya, Fiber Pathways of the Brain (Oxford University Press, 2006; 10.1093/acprof:oso/9780195104233.001.0001).

110. Brent A Vogt (ed.), Cingulate Neurobiology and Disease (Oxford University Press, 2009; 10.1093/oso/9780198566960.001.0001).

111. K. L. Bryant, L. Li, N. Eichert, R. B. Mars, A comprehensive atlas of white matter tracts in the chimpanzee. PLoS Biol 18, e3000971- (2021).

112. M. Catani, M. Thiebaut de Schotten, A diffusion tensor imaging tractography atlas for virtual in vivo dissections. Cortex 44, 1105–1132 (2008).

113. S. Walbridge, G. J. A. Murad, J. D. Heiss, E. H. Oldfield, R. R. Lonser, Technique for enhanced accuracy and reliability in non-human primate stereotaxy. J Neurosci Methods 156, 310–313 (2006).

114. R. W. Cox, AFNI: software for analysis and visualization of functional magnetic resonance neuroimages. Comput Biomed Res 29, 162–173 (1996).

115. B. Jung, P. A. Taylor, J. Seidlitz, C. Sponheim, P. Perkins, L. G. Ungerleider, D. Glen, A. Messinger, A comprehensive macaque fMRI pipeline and hierarchical atlas. Neuroimage 235, 117997 (2021).

116. K. J. Gorgolewski, T. Auer, V. D. Calhoun, R. C. Craddock, S. Das, E. P. Duff, G. Flandin, S. S. Ghosh, T. Glatard, Y. O. Halchenko, D. A. Handwerker, M. Hanke, D. Keator, X. Li, Z. Michael, C. Maumet, B. N. Nichols, T. E. Nichols, J. Pellman, J.-B. Poline, A. Rokem, G. Schaefer, V. Sochat, W. Triplett, J. A. Turner, G. Varoquaux, R. A. Poldrack, The brain imaging data structure, a format for organizing and describing outputs of neuroimaging experiments. Sci Data 3, 160044 (2016).

117. X. Wang, X.-H. Li, J. W. Cho, B. E. Russ, N. Rajamani, A. Omelchenko, L. Ai, A. Korchmaros, S. Sawiak, R. A. Benn, P. Garcia-Saldivar, Z. Wang, N. H. Kalin, C. E. Schroeder, R. C. Craddock, A. S. Fox, A. C. Evans, A. Messinger, M. P. Milham, T. Xu, U-net model for brain extraction: Trained on humans for transfer to non-human primates. Neuroimage 235, 118001 (2021).

118. J. Seidlitz, C. Sponheim, D. Glen, F. Q. Ye, K. S. Saleem, D. A. Leopold, L. Ungerleider, A. Messinger, A population MRI brain template and analysis tools for the macaque. Neuroimage 170, 121–131 (2017).

119. K. S. Saleem, A. V Avram, D. Glen, C. C.-C. Yen, F. Q. Ye, M. Komlosh, P. J. Basser, High-resolution mapping and digital atlas of subcortical regions in the macaque monkey based on matched MAP-MRI and histology. Neuroimage 245, 118759 (2021).

120. R. Hartig, D. Glen, B. Jung, N. K. Logothetis, G. Paxinos, E. A. Garza-Villarreal, A. Messinger, H. C. Evrard, The Subcortical Atlas of the Rhesus Macaque (SARM) for neuroimaging. Neuroimage 235, 117996 (2021).

